# Correlative Ultrastructural Mapping of Lewy Pathology Reveals Regional Diversity in Parkinson’s and Dementia with Lewy bodies

**DOI:** 10.64898/2025.12.19.695426

**Authors:** Notash Shafiei, Daria Proniakova, Marija Simjanoska, Anniken Mathea Rafnum Sjødal, Daniel Stähli, Lukas van den Heuvel, Marta di Fabrizio, Eve Aaron, Sandor Kasas, Mathis Solal Krause, Mallory Wittwer, Julika Radecke, Henning Stahlberg, Wilma DJ van de Berg, Amanda J Lewis

## Abstract

Lewy body diseases, including Parkinson’s disease (PD) and dementia with Lewy bodies (DLB), are defined by neuronal accumulation of misfolded α-synuclein (α-Syn), yet the ultrastructural diversity of these inclusions across brain regions and disease contexts remains unclear. Here, we applied large-scale correlative light and electron microscopy (CLEM) to map α-Syn pathology across cortical regions (entorhinal cortex, ENT; anterior cingulate cortex, AC; hippocampal CA2 region) and substantia nigra (SN) in clinically and pathologically confirmed PD and DLB donors. We identified pronounced regional heterogeneity in Lewy pathology, with cortical inclusions showing diverse maturation stages at the ultrastructural level, ranging from low-density fibrils interspersed with organelles to highly compact fibrillar inclusions. In the SN of DLB donors, we observed the full range of classical nigral LB morphologies previously described in PD. We additionally characterized diverse neuritic α-Syn pathologies in DLB and identified a distinct population of electron-dense, degenerating, α-Syn-positive cortical neurons not previously reported. Importantly, we found no significant difference in LB ultrastructure between PD and DLB in either cortical or nigral pathology. In contrast, quantitative analysis of >10,000 mitochondria revealed disease- and region-specific signatures of altered mitochondrial homeostasis. PD showed increased mitochondrial density and enlargement in the SN, whereas DLB showed increased mitochondrial density only in the ENT. Mitochondrial enlargement was exclusive to PD. These findings indicate that LB ultrastructure alone does not distinguish PD from DLB; instead, region-specific mitochondrial phenotypes may better reflect disease identity and regional susceptibility. Overall, we provide a high-resolution framework for human Lewy pathology in PD and DLB, revealing that ultrastructural responses to α-Syn pathology are driven primarily by neuronal identity and regional vulnerability. Our results highlight the need for disease- and region-specific models that capture human phenotypes to advance mechanistic understanding and therapeutic targeting of synucleinopathies.

## Introduction

Parkinson’s disease (PD) and dementia with Lewy bodies (DLB) are neurodegenerative disorders characterized by the abnormal neuronal accumulation of alpha-synuclein (α-Syn), forming Lewy bodies (LB) and Lewy neurites (LN)[9, 55]. Clinically, PD is defined by motor symptoms due to dopaminergic degeneration in the substantia nigra (SN), with cognitive impairment typically emerging later as Parkinson’s disease dementia (PDD)[1, 32, 54]. In contrast, DLB is diagnosed when dementia occurs prior to or within one year of the onset of parkinsonism[39, 67]. Despite these diagnostic distinctions, there is substantial overlap in clinical features and underlying pathology, leading to ongoing debate about whether they represent separate diseases or a spectrum of a shared synucleinopathy[15, 39, 46, 53].

Neuropathologically, the presence of Lewy pathology, particularly in the midbrain and limbic brain regions, together with the degeneration of dopaminergic neurons in the SN, represents the neuropathological hallmark of PD. In DLB, significant dopaminergic neuronal loss is also observed, but widespread cortical LB pathology is particularly prominent. Most PD patients progressively develop dementia over time (PDD) and, at end-stage, also show diffuse cortical Lewy pathology, making it difficult to distinguish neuropathologically from DLB[5, 54].

The ultrastructure of nigral inclusions in PD has been characterised in detail through numerous electron microscopy studies[6, 14, 18, 19, 31, 34, 51, 65]. Classical nigral LBs are described as spherical structures (5-25µm in diameter) composed of α-Syn fibrils interspersed with granular and vesicular components and arranged around an electron dense core, with immature “pale bodies” showing less organization[12, 34, 59, 63]. The fibrillar density and degree of structural organization within LBs have been linked to their stage of maturation[18, 34]. LNs, elongated α-Syn immunopositive inclusions within neuronal processes, are classically described to share a similar fibrillar composition[4, 51, 55]. However, more recent EM studies revealed a more heterogeneous spectrum of ultrastructures, with densely accumulated membranous material (vesicles, membrane fragments, cellular organelles) alongside α-Syn fibrils. Notably, some compact neuritic inclusions were found to consist almost entirely of membranous material with little to no fibrils[34, 51].

In contrast, the ultrastructure of cortical LB pathology in PD and DLB remains poorly understood. Immunohistochemistry studies describe morphological differences between brainstem and cortical LBs[6, 25, 30, 43, 45], and the limited electron microscopy studies report that cortical LBs are either granular, or generally fibrillar and loosely organized, but notably lacking the peripheral radiating fibrillar halo and electron dense core characteristic of brainstem LBs[20, 23, 27]. However, systematic regional comparisons have not been performed, hindering interpretation of disease-specific mechanisms and limiting the development of human-relevant models that recapitulate cortical involvement in synucleinopathies.

In this study, we used correlative light and electron microscopy (CLEM) to investigate the regional diversity of α-Syn pathology in PD and DLB. We compared α-Syn inclusions in the entorhinal cortex (ENT), anterior cingulate cortex (ACC), and examined pathology in the hippocampus (CA2) and SN of DLB donors, also describing neuritic pathology across multiple cortical regions in DLB for the first time. We show that LB ultrastructure varies by brain region, and not disease identity, with the ENT exhibiting a spectrum of fibrillar densities and the ACC showing only uniformly compact inclusions. Integrating this ultrastructural analysis with large-scale mitochondrial profiling, we also uncovered regional and disease-specific differences in mitochondrial morphology associated with α-Syn inclusions. These findings highlight how neuronal identity and the local pathological context shape the cellular response to α-Syn aggregation. Together, our findings extend previous ultrastructural descriptions of cortical Lewy pathology, providing new insight into its regional heterogeneity and molecular complexity in PD and DLB.

## Material and Methods

### Brain samples

We included four Dementia with Lewy Body, three Parkinson’s, one Alzheimer and two non-neurological control brain donors, who participated in the brain donation program from the Netherlands Brain Bank (www.brainbank.nl) with a post-mortem delay of <6 hours (Supplementary Table 1). Brain regions were dissected at autopsy according to the standardized procedure of the NBB (www.brainbank.nl)). Tissue blocks from the substantia nigra, and cingulate gyrus, entorhinal cortex and the CA2 region of the hippocampus used for CLEM were fixed at autopsy in 4 % formalin and 0.1 % glutaraldehyde in 0.15 M cacodylate buffer, supplemented with 2 mM calcium chloride, pH 7.4 for 24 hours. Next, tissue was stored in cacodylate buffer with 0.1% formalin until further processing.

For pathological diagnosis, seven µm-thick FFPE-embedded sections were immuno-stained using antibodies targeting α-Syn (clone KM51, 1:500, Monosan Xtra, The Netherlands), amyloid-β (clone 4G8, 1:8000, Biolegend, USA) and phosphorylated tau (p-tau, clone AT8, 1:500, Thermo Fisher Scientific, USA), following previously described protocol[38]. α-Syn pathology was staged according to Braak and McKeith criteria using BrainNet Europe (BNE) guideline[3]. Based on Thal amyloid-β phases scored on the medial temporal lobe[2], Braak neurofibrillary stages[62] and CERAD neuritic plaque scores[61], levels of AD pathology were determined according to NIA-AA consensus criteria[21]. Additionally, Thal CAA stages[60], presence of aging-related tau astrogliopathy (ARTAG)[29], microvascular lesions and hippocampal sclerosis were assessed.

### CLEM

Correlative light and electron microscopy (CLEM) was performed as described previously[50]. To localize pathologies of interest, free-floating vibratome slices were immunolabelled with phosphorylated α-Syn (pS129 Abcam ab51253; 1/1000 dilution) and neurofilament H (Sigma Ab5539; 1/500 dilution) incubated overnight at 4 °C and visualized with a donkey anti-rabbit Alexa 488, goat anti-chicken Alexa 558, and DAPI (Biolegend #422801; 1/800 dilution), incubated at room-temperature for 30 mins. Immuno-positive neurons were localised in the sections and imaged using a Leica Thunder microscope equipped with a K8 fluorescent camera, before being resin embedded. The region containing the immuno-positive neurons were excised by laser-capture microdissection (Leica LMD7; 5X objective, laser power-60, aperture 1, speed-5, specimen balance-0, pulse frequency 3500). Ultra-thin sectioning on excised regions was carried out on a UC7 ultramicrotome (Leica), with sections collected alternatively on glass slides (150-200 nm) and EM grids (80-150 nm). For immunohistochemistry a primary antibody against α-Syn (Antibody Clone 42, BD Biosciences # 610786; 1/100 dilution for 4 hours at room-temperature) with a secondary antibody (ImmPRESS Reagent Anti-Mouse Ig, Vector Laboratories) for 30 mins at room temperature were used. Antigen retrieval for α-Syn was carried out with 100 % formic acid for 10 minutes followed by steaming in Tris-EDTA, pH 9 for 30 mins at 100 °C.

### Imaging

Light microscopy images of selected glass slides containing immuno-positive regions were collected using a Leica Thunder microscope equipped with a DMI8 color camera. The entire section was imaged in overlapping tiles at 400 X or 630 X (oil immersion) magnifications, and image tiles were merged into a single image using the LAS X software (Leica Microsystems).

The fluorescent images were recorded with either a 40 X (HC PL FLUOTAR, NA 0.13), 100 X (HC PL APO, NA 0.45), or 200 X (HC PL APO, NA 0.8) air objective for overview images. Higher magnification images were recorded at 400 X (HC PL APO, NA 0.95), air objective or 630 X (HC PL APO CS2, NA 1.4) oil objective and z-stacks with a 0.3 μm step size. The light source for the blue channel has excitation and emission ranges of 395–440 nm; for the green channel, 480–520 nm; for the yellow channel, 555–590 nm; and for the red channel, 640–700 nm.

TEM images of electron microscopy grids consecutive to those imaged by light microscopy were collected at room temperature on a 120 kV Tecnai G2 Spirit TEM microscope operated at 80 kV with a LaB6 filament and a side-mounted EMSIS Veleta camera, or a CM100 Biotwin (Philips) operated at 80 kV with a Lab6 filament and bottom mount TVIPS F416 camera or a 120 kV TFS Biotalos with a LaB6 filament CETA camera or a 200 kV TFS Talos with a LaB6 filament CETA camera.

Electron and light microscopy images were adjusted for brightness and contrast using FIJI[44].

### Mitochondria Segmentation

43 EM images were manually annotated as a training dataset using Napari[48]. The images were from two brain regions: the ENT and SN. These images varied in size and were acquired at three different magnifications to ensure robustness and generalizability of the model. Binary masks were processed in Napari then converted into polygonal annotations compatible with QuPath[7] v0.5.0 enabling streamlined visualization and data management. The dataset was split into a training and evaluation set: 30 images were randomly selected for training, and the remaining 13 were used for testing and performance assessment. We fine-tuned the Cellpose Cyto2 model[40, 41], an architecture that predicts both probability maps and spatial flow fields from each pixel to object centers, to guide instance reconstruction using the built-in deep learning extension of QuPath. The model was trained for 75 epochs on the prepared training set. Training was conducted using a single NVIDIA GeForce RTX 3060 GPU. To assess segmentation performance, we computed the Intersection over Union (IoU) metric on a per-object basis. A linear sum assignment algorithm (Hungarian method) was used to optimally match predicted and ground-truth objects after converting QuPath detections into label masks. This object-level evaluation ensured a robust measure of segmentation accuracy, accounting for both shape and location. For higher accuracy, the output segmented images were proofread using segment anything (SAM)[24].

### Statistical analysis

To assess the number of mitochondria per cell area, we first counted the number of segmented mitochondria in each cell. We then calculated the cell area by multiplying the number of pixels per cell by the surface area of the square pixels for each image. Mitochondrial density was determined by dividing the number of mitochondria (number of segmented mitochondria) by the corresponding cell area. To measure average mitochondrial size per cell, we calculated the area of each mitochondrion by counting its pixels and multiplying by the squared pixel size. The average mitochondrial area was then computed for each cell. All calculations were performed using custom Python codes, available on GitHub. Mitochondrial size measurements were analyzed using linear mixed-effects models to account for the nested experimental design of the study. Because multiple cellular measurements were obtained from each patient, patient identity was included as a random effect to account for the non-independence of repeated measurements originating from the same individual.

For comparisons between α-synuclein-positive and α-synuclein-negative cells within the same disease group, cell type was included as a fixed effect, while patient identity was modelled as a random intercept. Separate models were fitted for positive and negative cell populations where appropriate. Comparisons between disease-associated cell populations and control samples were performed using analogous mixed-effects models.

Model parameters were estimated using restricted maximum likelihood (REML). Fixed-effect coefficients and associated p-values were extracted from each model. Statistical significance was defined as (p < 0.05). Graphs were made in GraphPad Prism version 10.5.0.

## Results

To assess how α-Syn inclusion ultrastructure varies across brain regions and synucleinopathies, we examined the ultrastructure of cortical α-Syn pathology in the ACC and ENT from clinically and pathologically confirmed PD and DLB donors (Supplementary Table 1). The cohort consists of three DLB cases, five PD cases, and one mixed-pathology case that was clinically diagnosed as DLB but pathologically classified to have more AD pathology than typically observed (DLB+AD).

LBs were identified within these cases by their immunofluorescence for α-Syn phosphorylated on serine 129 in free-floating brain sections (EP1536Y, Abcam Ltd., Cambridge, UK) or by post-embedding immunohistochemistry on ultra-thin sections (clone 42, BD Biosciences), consistent with methods established previously[49]. Intraneuronal localization of LBs (*i.e*., somatic *vs*. neuritic) was confirmed by the presence of a nucleus and/or other cellular features identified by their ultrastructural morphology such as cell organelles, including the Golgi apparatus or the presence of lipofuscin. In total, from the DLB cases we correlated 46 LBs and 12 LNs in the ENT, 24 LBs in the ACC, 2 LBs and 6 LN in the CA2, and 70 LBs and 40 LNs in the SN. From the PD donors we correlated 16 LBs in the ENT and 11 LBs in the ACC (Supplementary Table 2). LBs in the SN of PD cases were not described here as the same cases have been comprehensively characterised in our recent parallel study[34]. LNs in the ACC of PD and DLB cases, as well as in the ENT and CA2 of PD cases, were not abundant, which limited the number of images we could obtain to characterize their ultrastructure (Supplementary Figure 1 and 2). Representative images of each ultrastructure are shown here, the full dataset is available at the BioImage Archive.

### Ultrastructural diversity of cortical Lewy bodies in DLB and PD

In the ENT of both PD and DLB, LBs displayed a predominantly fibrillar composition, ranging in diameter from 8 to 11 µm, from which we identified three distinct ultrastructural morphologies (Figure 1). The first consisted of a mixture of organelles such as mitochondria, lysosomes and other vesicular components, interspersed with a low-density of disorganized fibrils (Figure 1a, d). The second morphology featured disorganized fibrils mixed with lipids and vesicles, but with a higher fibrillar density compared to the first morphology. Most mitochondria within these inclusions were located at the LB periphery, excluded from the central fibrillar region (Figure 1b, e). The third morphology consisted of a dense and compact fibrillar ultrastructure, lacking any intermixed organelles (Figure 1c, f). This morphology was identified twice amongst the DLB cases but was observed more frequently in the PD donors (Supplementary Table 2). These inclusions consistently displayed a densely packed fibrillar ultrastructure across the different CLEM cycles. In these cells, the nucleus was typically located at the cell periphery, in contrast to aSyn-negative neurons showing a central nucleus (Supplementary Figure 3), and there was noticeable empty space between the neuron and the surrounding tissue.

**Figure 1.**
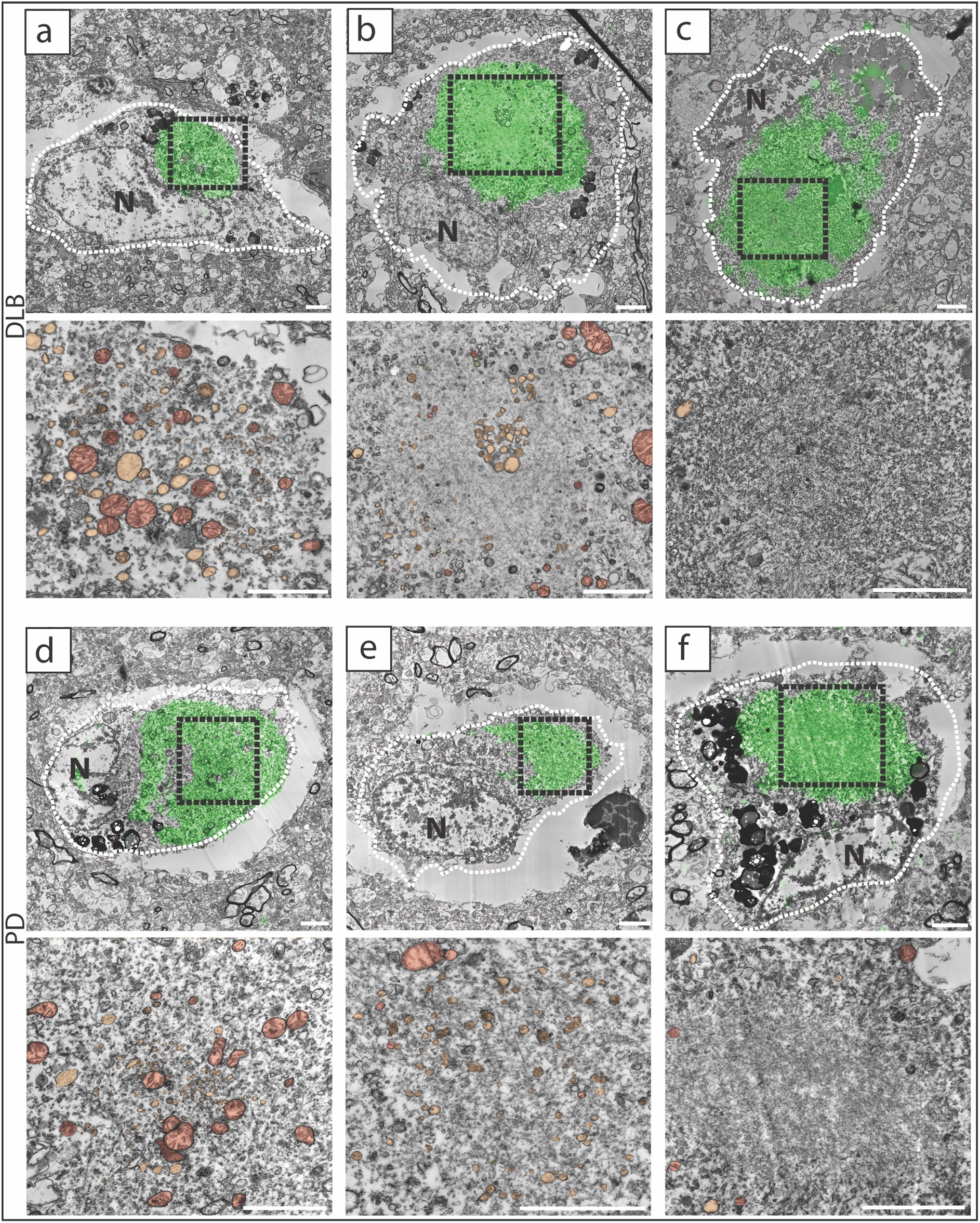
Ultrastructural diversity of LBs in the ENT of DLB and PD. Representative CLEM images of three distinct LB morphologies identified in ENT neurons. (a, d) Low-density fibrils interspersed with mitochondria, vesicles and other organelles. (b, e) Intermediate-density fibrils intermixed with vesicles. Mitochondria are not observed within the fibrillar core but localized to the periphery. (c, f) High density compact fibrillar inclusions lacking other organelles and vesicles. In all cases, the nucleus (N) is displaced to the cell periphery, with visible separation between the neuron and surrounding tissue. In all panels, dashed lines outline neuronal boundaries; black dashed boxes indicate higher magnification views. Mitochondria (red) and other organelles (yellow) are annotated. The IHC α-Syn immunostaining (green) is overlaid onto the EM micrograph. Scale bars: 2 µm.

In the ACC for both PD and DLB, LBs ranged in diameter from 7 to 10 µm and consistently showed a compact, densely packed ultrastructure composed of disorganized fibrils and devoid of any intermixed organelles (Figure 2), in agreement with previous reports[20, 27, 28]. Cellular organelles, such as mitochondria and vesicles, were located at the periphery of the compact fibrils. Like the observations in the ENT, there was separation between the inclusion and the nucleus filled with cellular material, and empty space was often observed between the cell membrane and surrounding tissue.

**Figure 2.**
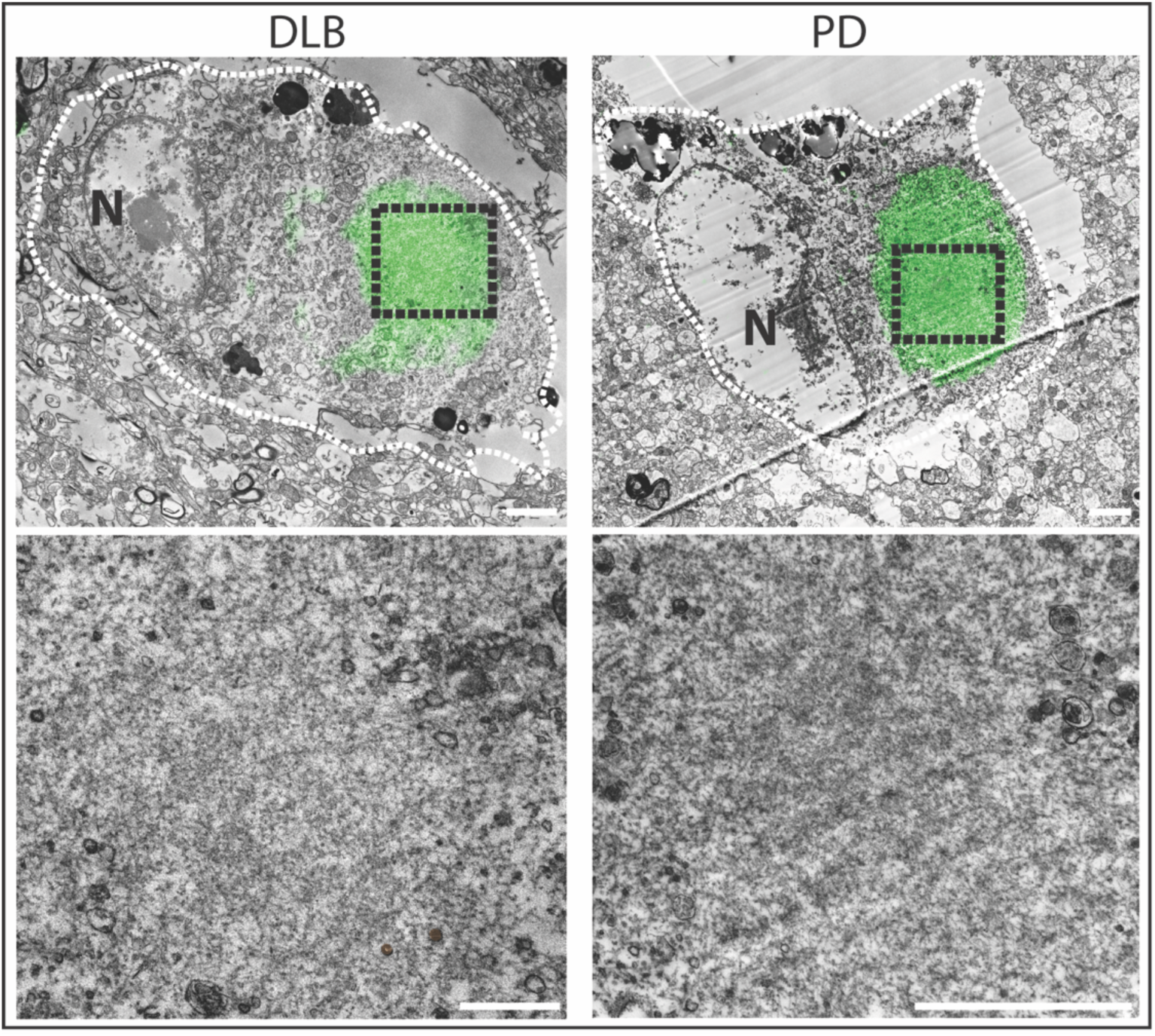
Compact mature fibrillar LBs predominate in ACC neurons. CLEM images showing uniformly dense fibrillar LB ultrastructure, devoid of internal organelles, in the ACC of PD and DLB donors. In all cases, the nucleus (N) is displaced to the cell periphery, with visible separation between the neuron and surrounding tissue. In all panels, dashed lines outline neuronal boundaries; black dashed boxes indicate higher magnification views. The IHC α-Syn immunostaining (green) is overlaid onto the EM micrograph. Scale bars: 2 µm.

In the hippocampus CA2 region of DLB cases, rare somatic inclusions were observed. We could only observe a low density of disorganized fibrils, mixed with organelles like vesicles and mitochondria which resembled the ultrastructure observed in the ENT (Supplementary Figure 4).

Ultrastructural analysis of LBs in the SN revealed morphologies previously reported for PD[6, 14, 18, 19, 34, 65], but not yet described for DLB (Figure 3). These morphologies included pale bodies composed of diffuse and disorganized fibrils intermixed with clustered mitochondria (Figure 3a), one-layer LBs comprising disorganized fibrils with peripheral organelles (Figure 3b), two-layer LBs containing a dense fibrillar core surrounded by a layer of disorganized fibrils (Figure 3c, Supplementary Figure 5), and a single observation of a classical three-layer LB with an electron dense core, a middle layer of densely packed fibrils, and an outer layer containing radiating fibrils (Figure 3d). It was unclear whether the three-layer LB was intracellular or extracellular, as no nucleus or other cellular features could be associated with the inclusion, even after tracking it on the adjacent sections through the volume (Supplementary Figure 6). In addition to the peripheral mitochondria that frequently surround the LBs (as we previously reported as a feature of LBs in PD[34]) we also observed clusters of mitochondria inside autophagosomes in the vicinity of the two-layer LB (Figure 3c). We also observed many instances of single neurons containing multiple LBs, another hallmark commonly seen in the SN in PD[34, 55, 64] cases (Supplementary Figure 7).

**Figure 3.**
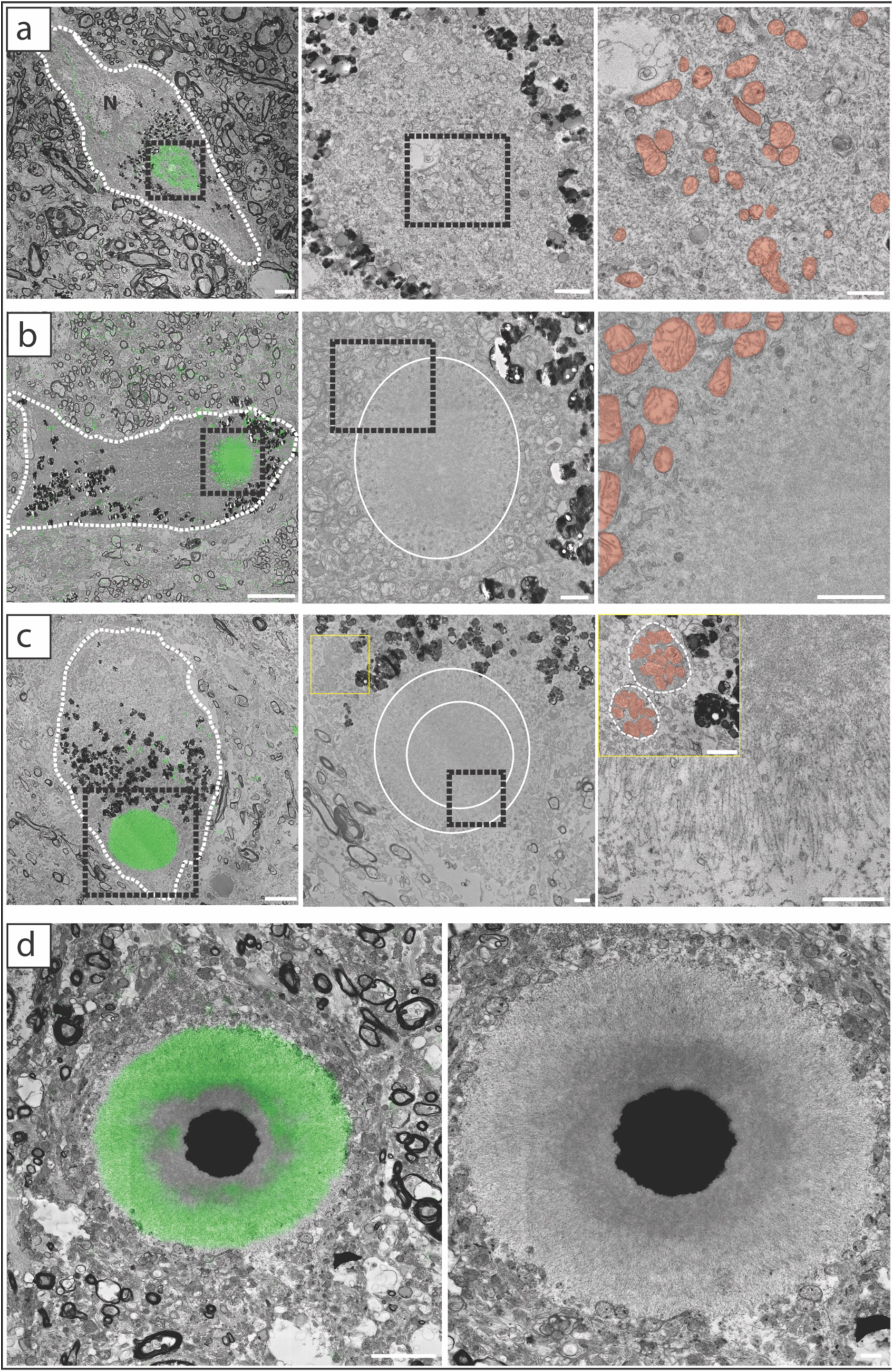
Classical nigral LB ultrastructures in DLB. CLEM images showing the ultrastructural categories identified in SN neurons from DLB donors, analogous to PD pathology. a) a pale body with low-density fibrils intermixed with clustered mitochondria. b) a one-layer LB, with disorganised fibrils. Clustered mitochondria are observed surrounding the inclusion. c) a two-layer LB, with disorganised fibrils in the centre and a linear arrangement of fibrils on the periphery. Autophagosomes containing mitochondria were observed in the vicinity of the LB (yellow box). d) a three-layer LB, with an electron dense core, dense fibrils in the middle layer and a linear arrangement of fibrils in the outer-most layer. In all panels, dashed lines outline neuronal boundaries; black boxed areas indicate higher magnification views. The IHC α-Syn immunostaining (green) is overlaid onto the EM micrograph. The nucleus (N) and mitochondria (red) are annotated. Scale bars (low mag): 5 µm, others 1 µm.

### Neuritic pathology shows heterogeneous ultrastructural morphologies

Neuritic pathology in the ENT and SN of DLB (Figure 4, Supplementary Figure 8), the ENT of PD cases (Supplementary Figure 2a), and in the CA2 in DLB (Supplementary Figure Supplementary Figure 9) and PD (Supplementary Figure 2b), displayed heterogeneous ultrastructural morphologies. These ultrastructures predominantly included membranous forms, occasionally mixed with very low-density fibrils (Fig 4a, d), disorganized fibrillar cores surrounded by other organelles (Figure 4b, e), or completely fibrillar neurites devoid of any organelle or membranous material (Figure 4c, f). Notably, three examples of fibrillar inclusions in the ENT of DLB cases were observed containing cores of material varying in electron density (Figure 4g, Supplementary Figure 10).

**Figure 4.**
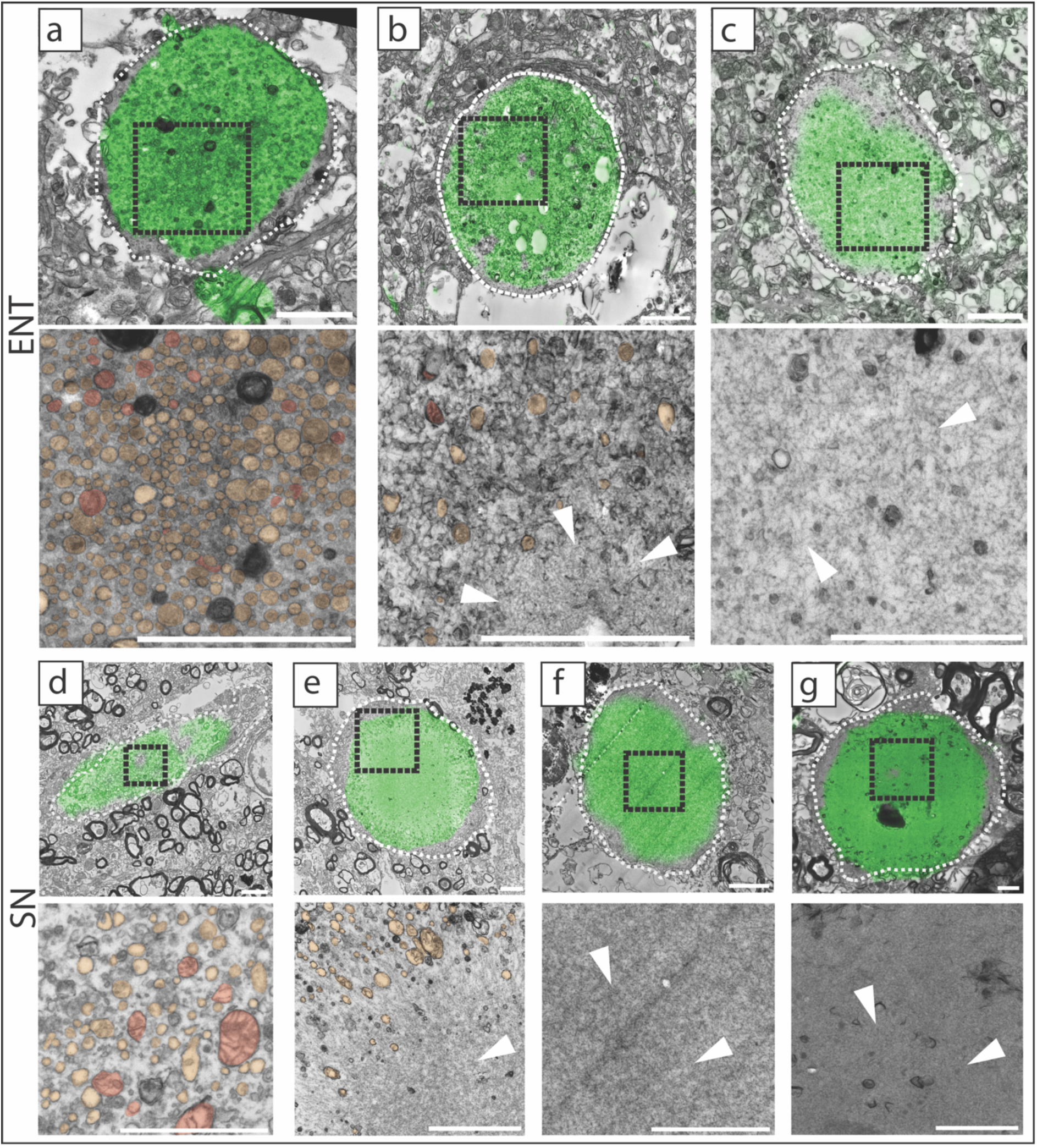
Neuritic pathology in DLB shows diverse fibrillar and membranous compositions. Representative CLEM images of LNs in the ENT (a-c) and SN (d-g). Three primary morphologies were observed: Membrane rich inclusions (a, d) inclusions containing a fibril-rich core surrounded by cellular organelles and vesicles (b, e) and fully fibrillar structures lacking internal organelles (c, f). g) Some fibrillar neurites contained an electron dense core. Panels a–c and e–g show cross-sections of neurites cut perpendicular to the imaging plane, whereas panel d shows a longitudinal view of a neurite extending parallel to the imaging plane. Further examples showing longitudinal views of neuritic pathology are shown in Supplementary Figure 8. In all panels, dashed lines outline neuritic boundaries; black boxed areas indicate higher magnification views. The IHC α-Syn immunostaining (green) is overlaid onto the EM micrograph. Mitochondria (red), other vesicles and organelles (yellow) and regions of fibrils (white arrowheads) are annotated. Scale bars: 1 µm.

### Other neuronal cortical pathology

In the cortex of both PD and DLB, we identified a population of α-Syn positive cells with distinct morphological features not previously described. These cells were characterized by an electron dense cytoplasm, giving them a dark appearance, and containing vacuoles and lipid droplets or lipofuscin granules (Figure 5). The nucleus in these cells also appeared dark, was irregularly shaped, and was often displaced into the somal processes. No comparable ultrastructures were observed in α-Syn immunonegative cells within the same areas, or within the healthy control tissue, although the possibility that such cells may be present in unexamined portions of the tissue cannot be excluded.

**Figure 5.**
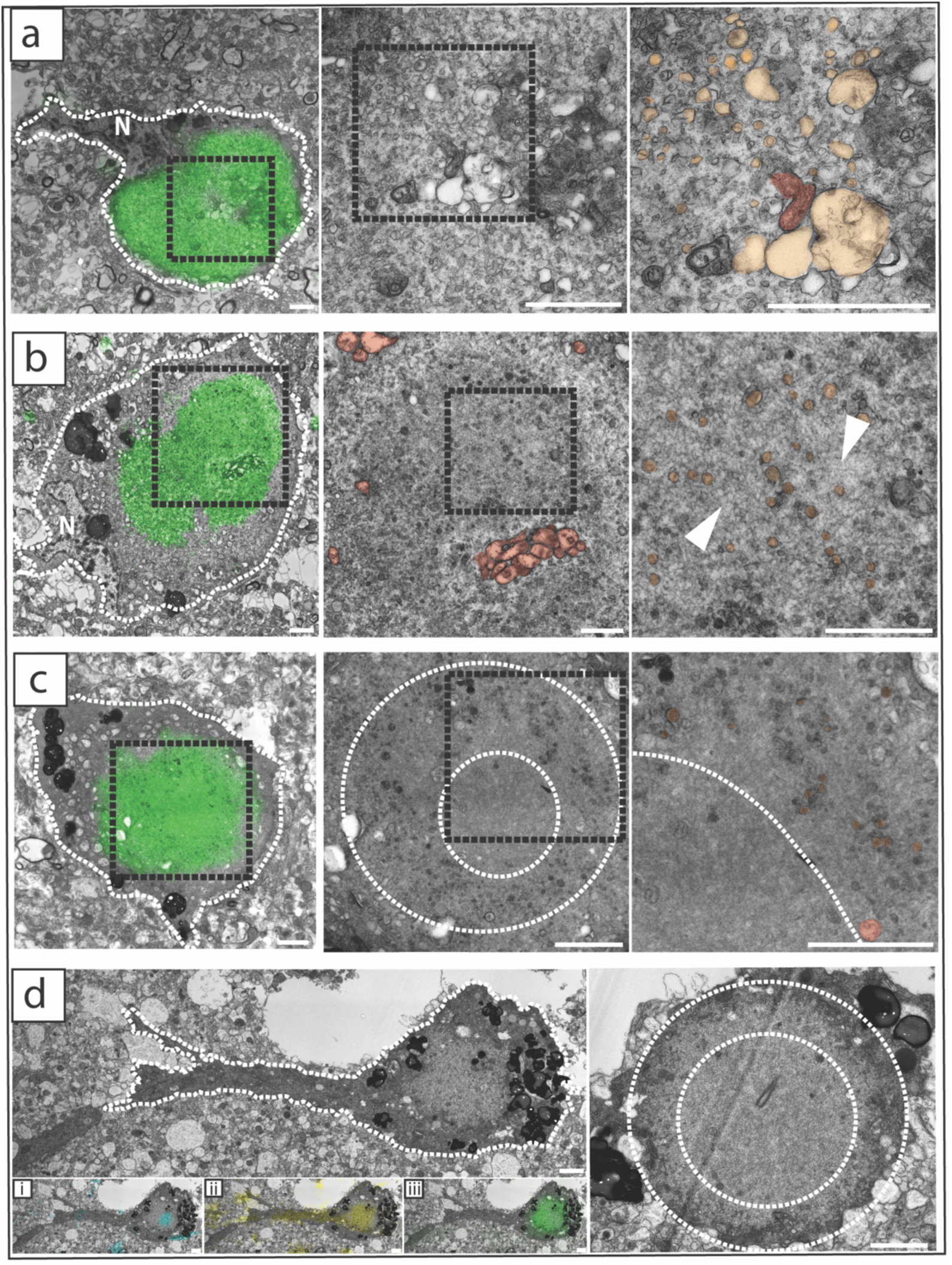
Electron dense degenerating α-Syn neurons in the cortex. Electron-dense neuronal cells containing α-Syn pathology in DLB ENT (a, b) and ACC in DLB (c) or PD (d). In all panels, dashed lines outline neuritic boundaries; black boxed areas indicate higher magnification views. (A, B) ENT neurons show either low-density or higher-density fibrils with clustered mitochondria. (C, D) ACC examples show layered fibrillar organization resembling staged LB maturation. (a-c) The IHC α-Syn immunostaining (green) is overlaid onto the EM micrograph. (d) The α-Syn immunofluorescence (green), DAPI (blue) and co-localization with neurofilament-H immunofluorescence (yellow) is overlaid onto the EM micrograph, confirming neuronal identity. Mitochondria (red), other vesicles and organelles (yellow) and regions of fibrils (white arrowheads) are annotated. Scale bars: 2 µm.

These electron dense pathological cells were observed most frequently in the ENT of DLB cases, showing ultrastructures that either lacked clearly visible fibrils amongst accumulated membranous material, including disrupted mitochondria, membrane fragments, and vesicles, in the core (Figure 5a), or with a higher density of fibrils intermixed with vesicles, and clustered mitochondria at the periphery (Figure 5b).

In the same region, we also observed a second type of α-Syn-positive pathological cell that differed markedly from the two morphologies described above. Although it similarly showed a very dense and dark cytoplasm, its α-Syn immunostaining pattern was discontinuous and patchy, rather than forming a single coherent inclusion. Ultrastructurally, this cell contained densely packed fibrils that were uniformly oriented in a regular, aligned pattern, distinct from the randomly arranged fibrils seen in the other electron-dense inclusions. Unlike the other similar cells in this region, this cell also lacked any clearly identifiable somal processes. (Supplementary Figure 11).

In the ACC, individual examples of electron dense α-Syn-positive cells were observed in one PD and one DLB case (Figure 5c, d). These inclusions exhibited a layered architecture, consisting of a dense fibrillar core surrounded by a peripheral layer of lower-density fibrils intermixed with vesicles and membrane fragments. Although this layering resembled that of two-layer LBs in the SN, no ordered or radiating fibrillar arrangements were detected. In the PD example, a cross section through the entire cell showed an electron-dense cell soma and similarly electron-dense neuritic process extending from it. Correlation of this cell with the pre-embedding fluorescence imaging showed strong neurofilament heavy chain (NF-H) immunoreactivity (Figure 5d), confirming a neuronal identity of these α-Syn-positive cells. In this example, no nucleus was detected either within the 40 μm immunofluorescence volume or across multiple CLEM imaging cycles following the inclusion. The absence of a visible nucleus, together with the pronounced electron density and presence of vacuoles observed in other inclusions of this cell type, suggests that these neurons may have been actively undergoing degeneration.

### Mitochondrial characteristics and mechanisms in DLB and PD

During our regional analysis of the various LB ultrastructures, we noted apparent differences in the mitochondrial morphology between cells containing α-Syn inclusions in the cortex and those in the nigra. To assess this systematically, we segmented over 10,000 mitochondria from more than 115 CLEM images obtained from the SN and ENT regions from two DLB, one DLB+AD, three PD and two non-neurological control cases. The ACC was excluded from this analysis due to the lack of non-neurological control tissue from this region. For each neuron, the mitochondria were segmented and quantified for average size and mitochondrial density per cell (Figure 6; examples of segmentations shown in Supplementary Figures 3, 12 and 13). As multiple cells were sampled from individual donors, mitochondrial measurements were averaged per cell prior to analysis, and the resulting cell-level analyses were used to assess the heterogeneity of mitochondrial phenotypes within pathological populations. Analysis included LB-containing neurons in cortical regions, LB and pale body containing neurons in the SN (α-Syn^+^), adjacent neurons immuno-negative for α-Syn (α-Syn^-^), and α-Syn immuno-negative neurons from the control donors (C). The DLB + AD case was excluded from the analysis to avoid any confounding effects due to its mixed pathology.

**Figure 6.**
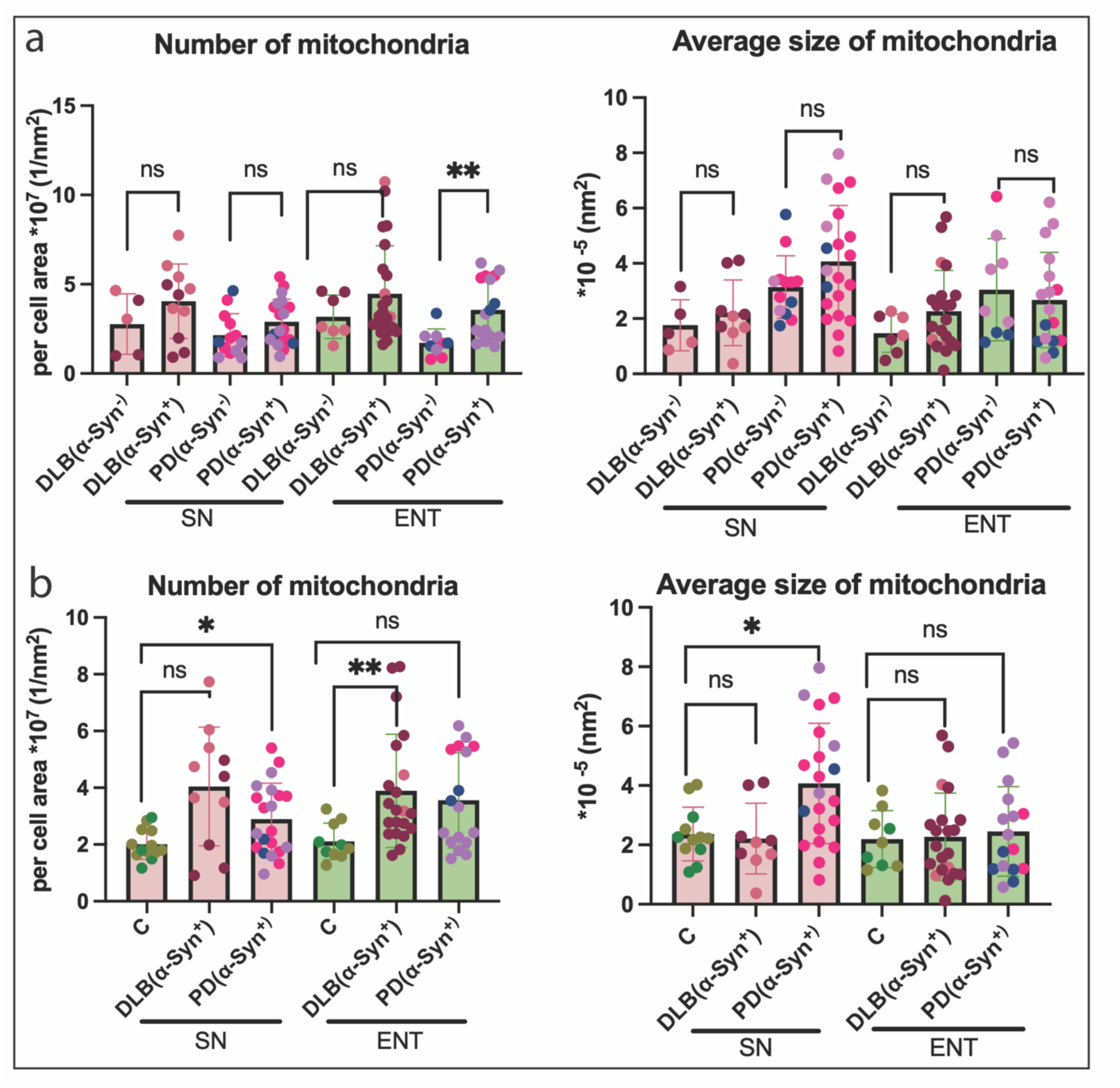
Mitochondrial alterations associated with α-Syn pathology. a) Within disease comparisons of mitochondrial density (number) and average size in α-Syn- and α-Syn+ neurons from PD and DLB cases. In PD, α-Syn pathology was associated with increased mitochondrial density in the ENT with no significant changes in the mitochondrial size for either brain region or disease. b) Disease vs control comparisons: In the SN, PD neurons showed increased mitochondrial density and size compared to controls, whereas DLB showed no significant changes. In the ENT, mitochondrial density was increased in DLB neurons compared to controls, while mitochondrial size was unchanged in both diseases. Each data point represents one cell, coloured by donor. Statistical significance was calculated using a mixed linear model and indicated as follows: p < 0.05 (*) and p < 0.01 (**). Outliers (not shown) were excluded from the p-value calculations. The average size of mitochondria is divided by 10 ^(5)^ and the number of mitochondria per cell are multiplied by 10^(7)^.

To determine whether mitochondrial alterations were specifically associated with α-Syn inclusions within diseased tissue, we compared α-Syn^+^ neurons to adjacent α-Syn^-^ neurons in the same region (Figure 6a). No significant differences in mitochondrial density were observed in the SN of PD or DLB cases, or in the ENT of DLB cases. In contrast, the ENT of PD cases showed a significant increase in the mitochondrial density in α-Syn^+^ neurons compared to adjacent α-Syn^-^ neurons (p<0.01). As mitochondrial density was quantified across the entire cell, this change reflects a global alteration in mitochondrial abundance rather than local redistribution of mitochondria around α-Syn inclusions. No significant differences in mitochondrial size were observed between α-Syn^+^ and α-Syn^-^ neurons in either region or disease group. These results indicate that significant inclusion-associated mitochondrial alterations within diseased tissue were only detected in the ENT of PD cases.

We next evaluated mitochondrial alterations in α-Syn^+^ neurons from diseased brains relative to non-neurological controls (Figure 6b). In the SN of PD cases, both mitochondrial density and average size were significantly increased compared to controls (p < 0.01), whereas no significant changes were observed in the SN of DLB cases. <u>In</u> the ENT, mitochondrial density was significantly increased in DLB cases compared to controls (p < 0.05), while PD cases showed no significant difference compared to controls. The average mitochondrial size in the ENT was unchanged in both diseases. These results show distinct patterns of mitochondrial alterations across diseases and brain regions, relative to non-neurological controls, with the most pronounced changes observed in the SN of PD cases.

## Discussion

Distinguishing the mechanisms that drive pathology in PD and DLB remains challenging and the extent to which ultrastructural features differ by disease or brain region has not been systematically evaluated. To address this gap, we applied CLEM across multiple cortical regions and the SN in clinically and pathologically confirmed PD and DLB donors, integrating a detailed analysis of α-Syn inclusion morphology with quantitative mitochondrial profiling. We found that while Lewy body and neuritic ultrastructures varied within a regional context, they did not differ between PD and DLB, indicating that inclusion architecture alone cannot resolve the disease identity. Instead, altered mitochondrial phenotypes emerged as a distinguishing feature with distinct regional- and disease-specific differences in mitochondrial density and size.

### Regional Identity, Not Disease Type, Shapes Lewy Body Ultrastructure

PD and DLB showed indistinguishable α-Syn inclusion ultrastructures across all studied brain regions, indicating that disease identity does not determine LB ultrastructure. These findings support recent arguments that DLB and PDD may share substantial underlying biological mechanisms, despite differing clinical trajectories[16, 46]. For both diseases, the cortical regions displayed region-dependent variability: the ENT showed a variety of ultrastructures ranging from loosely organized fibril-organelle mixtures to compact fibrillar inclusions (Figure 1), whereas the ACC contained uniformly dense, fully compacted LBs (Figure 2). Earlier studies have reported the ultrastructure of cortical LBs to appear either granular or fibrillar and lacking the dense core and peripheral halo of radiating α-Syn filaments characteristic of brainstem LBs[20, 23, 28]. Our findings expand on these earlier descriptions, revealing greater ultrastructural heterogeneity than previously recognized. Notably, we did not observe the previously reported granular morphology. This difference may reflect improved ultrastructural preservation, as our samples were obtained with short post-mortem delays and processed using fixation protocols optimised for membrane and protein preservation. These conditions enhance overall ultrastructural retention compared with earlier preparation methods, in which loss of cellular components may have contributed to the appearance of more granular-like inclusions[33].

In the SN of DLB donors, we observed the full spectrum of classical LB morphologies, including pale bodies, one-, two-, and three-layer halo LBs (Figure 3), mirroring established features in PD[14, 19, 34, 51, 65]. This is, to our knowledge, the first detailed report of the complete range of classical nigral LB morphologies in DLB cases.

In both diseases we observed neuritic α-Syn pathology to be ultrastructurally diverse (Figure 4), including predominantly membranous inclusions to mixed membrane-fibril aggregates and fully fibrillar inclusions, also consistent with recent observations for the SN in PD[34]. In contrast to somal pathology, the neuritic inclusions did not display any obvious regional differences.

Taken together, these results indicate that regional neuronal identity strongly influences LB and LN morphology, whereas the underlying synucleinopathy (PD or DLB) does not. This is consistent with recent cryo-EM studies demonstrating that α-Syn fibrils from PD and DLB adopt the same core “Lewy fold”[68], indicating that fibril architecture alone does not account for clinical or neuropathological divergence between these disorders. Instead, factors such as regional neuronal identity, local microenvironment, and the temporal order in which brain circuits are affected, likely shape the cellular consequences of α-Syn aggregation[37, 56]. Although cryo-EM structures have not yet been systematically compared across brain regions within the same individual, and our EM resolution does not permit direct assessment of fibril polymorphs, our findings support emerging perspectives in which the context of aggregation, rather than fibril conformation alone, determines pathological outcomes[66].

### Regional ultrastructural heterogeneity in the cortex reflects distinct maturation trajectories

Previous ultrastructural studies of the LB maturation stage[18, 34] and *in vitro* models of α-Syn aggregation[35, 36] have proposed that fibrillar density reflects inclusion maturity, with loosely organized inclusions representing earlier stages and densely compact fibrillar structures marking later progression. Our data support this concept: in the ENT, which is among the earliest cortical regions affected in synucleinopathies, we observed a broad spectrum of fibrillar densities, consistent with inclusions undergoing progressive structural compaction. Notably, less densely packed morphologies were more frequent in DLB than PD, aligning with the typically shorter disease duration and accelerated cortical involvement in DLB.

In contrast to our observations in the ENT, LBs in the ACC consistently showed a compact and high-density fibrillar ultrastructure, with no evidence of intermediate or low-density fibrillar inclusions (Figure 2). The presence of exclusively mature LBs in this region is consistent with the fact that all cases in our cohort were end-stage (Braak 6) and therefore have had sufficient time to develop into fully compacted inclusions. However, according to Braak staging[8], the ENT is affected earlier than the ACC, and so the observed LB heterogeneity in the ENT, but not in the ACC, suggests region-specific differences in neuronal vulnerability and local cellular environments that influence LB maturation.

Consistent with recent models proposing that timing is a major determinant of pathological outcome in Lewy body diseases[37], our observations suggest that LB ultrastructure can reflect not only the temporal sequence of pathology spread, but also intrinsic neuronal and regional properties that modulate LB maturation (Figure 7). The analysed cohort in our study conforms to established Braak α-Syn staging. Therefore, our interpretation is aligned with this canonical progression model, and we acknowledge that this may not apply to atypical cases with divergent patterns of pathology.

**Figure 7.**
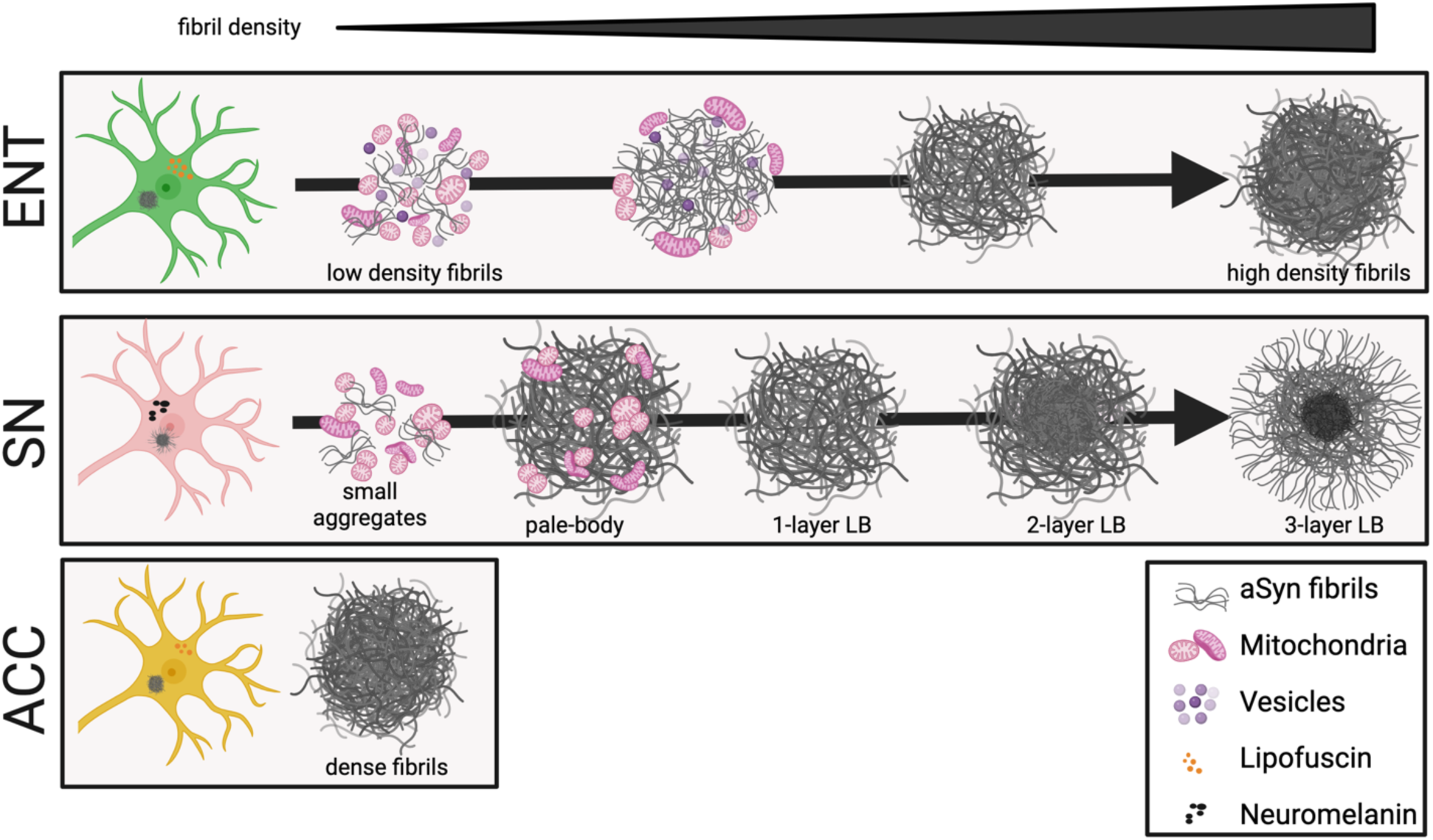
Proposed model for region-specific LBs maturation stages for PD and DLB. This model summarizes the distinct ultrastructural states of α-Syn inclusions observed across vulnerable brain regions based on the density and organization of fibrillar α-Syn structures identified by CLEM. In the ENT, α-Syn inclusions range from early, loosely organized fibril–organelle mixtures to compact fibrillar LBs, consistent with progressive maturation. In the SN, dopaminergic neurons exhibit the full sequence of classical LB morphologies, including pale bodies and one- to three-layer LBs. In the ACC, inclusions are uniformly compact and fibrillar, suggesting accelerated maturation in this late-affected region. This schematic integrates region-dependent neuronal identity and timing of pathological involvement, illustrating how local cellular environment shapes the structural outcomes of α-Syn aggregation in PD and DLB.

### Degenerating Neurons Reveal Additional Pathological Endpoints

An additional factor contributing to the regional differences in cortical α-Syn pathology is the presence of the electron-dense α-Syn-positive degenerating neurons, observed in both the ACC of PD and DLB cases, but not in the ENT for either disease (Figure 5). Increased electron density is a hallmark of advanced cellular degeneration, reflecting cytoplasmic and organelle condensation, protein aggregation, and accumulation of degradative bodies such as lysosomes and autophagosomes. Consistently, these neurons contained numerous vacuoles, and their nuclei exhibited condensation, fragmentation, shrinkage, chromatin clumping, or, in some cases, complete loss. The absence of similar morphologies in α-Syn–negative cells suggests that this degeneration is linked to α-Syn pathology.

The regional pattern we observed for these degenerating pathological neurons suggests an interplay between selective vulnerability and the temporal sequence of pathological involvement. The ENT is among the earliest cortical regions affected in synucleinopathies[3, 8]; therefore, neurons here experience a longer cumulative exposure to α-Syn aggregation. For DLB, where there is a more rapid and extensive progression of cortical pathology compared to PD[22, 47], this prolonged pathological burden is sufficient to trigger neuronal degeneration in both the ENT and ACC. In contrast, in PD, the ENT pathology may not accumulate sufficient pathological burden by end-stage disease to trigger this level of degeneration, consistent with the later and slower spread of cortical α-Syn pathology compared with DLB. However, once the ACC becomes affected, its inherent vulnerability means that both diseases reach a degenerative threshold, leading to the coexistence of mature and degenerating inclusions.

These degenerating pathological neurons were not observed in the SN for either disease, likely reflecting either the severe neuronal loss characteristic of this region by end-stage disease (∼50% at time of diagnosis[13, 26] or reflecting intrinsic differences in cellular vulnerability, with cortical neurons being less resilient to α-Syn aggregation than dopaminergic SN neurons.

Together, these data indicate that neuronal degeneration reflects both selective vulnerability and cumulative exposure to α-Syn pathology, with the susceptibility threshold reached more extensively in DLB than in PD.

### Region-specific mitochondrial alterations distinguish PD and DLB

Despite the similar ultrastructural appearance of LB pathology in PD and DLB, mitochondrial phenotypes differed between diseases and brain regions, consistent with previous reports [10, 11, 42, 52]. The clearest mitochondrial alterations were observed in the SN of PD cases, where both mitochondrial density and enlargement were increased relative to controls. In contrast, DLB cases did not show mitochondrial enlargement in the SN, while cortical regions primarily showed altered mitochondrial density without accompanying enlargement. These findings suggest that mitochondrial responses to α-Syn pathology differ across vulnerable neuronal populations and disease contexts.

Notably, mitochondrial enlargement was restricted to the SN of PD and was present not only in α-Syn⁺ neurons, but also in neighbouring α-Syn⁻ neurons. This suggests that mitochondrial dysfunction in PD extends beyond neurons containing detectable fibrillar α-Syn inclusions and may reflect broader disturbances in cellular homeostasis within vulnerable nigral circuits[17, 57, 58]. This interpretation is consistent with previous ultrastructural studies showing widespread organelle disruption surrounding nigral LBs, including degenerating mitochondria, autophagic vesicles, and ER–lysosomal abnormalities[51]. However, the absence of a similar phenotype in DLB further indicates that dopaminergic neuronal vulnerability alone is insufficient to account for these mitochondrial changes.

The increased mitochondrial density observed in cortical regions without accompanying enlargement may reflect a distinct form of mitochondrial remodelling compared to the overt structural abnormalities observed in the SN of PD. Together, these findings indicate that although PD and DLB share superficially similar LB ultrastructures, their associated cellular responses diverge substantially across brain regions.

While this study focused on mitochondrial alterations, additional organelles including lysosomes, ER, lipofuscin, and autophagic vesicles, may also display disease- or region-specific changes that warrant further quantitative assessment. Future studies integrating region-specific cryo-EM with ultrastructural and metabolic profiling will be important to determine how a common α-Syn fibril fold drives distinct cellular remodelling pathways across different neuronal populations.

### Limitations and Future Directions

Although CLEM is a powerful approach that provides high-resolution information and detailed ultrastructural context, the cohort size in our study was constrained by the tissue availability which may limit the breadth of regional comparisons. Additionally, CLEM sampling necessarily only covers a fraction of the affected regions in each donor, and the number of inclusions analysed per case was also affected by inter-patient variability in pathological burden. Therefore, the complete spectrum of regional heterogeneity may not have been fully captured in our dataset. Nonetheless, the consistency of key features across donors supports the robustness of our observations.

A key limitation of this study is that all analyses were performed in end-stage disease tissue, providing a cross-sectional view of pathology. In addition, donor-level statistical analyses were limited by the small number of biological replicates available for ultrastructural human brain studies. While mixed linear models were used to account for the nested structure of the data, the limited number of donors reduces statistical power and constrains robust estimation of donor-level variability and disease-associated effects. Larger donor cohorts will be required to determine whether the observed mitochondrial phenotypes represent reproducible disease-specific patterns.

As a result, we cannot directly determine temporal relationships, including whether mitochondrial alterations arise upstream or downstream of α-Syn aggregation. Nor can we confirm progression of individual inclusion morphologies. It remains possible that certain ultrastructural forms may persist rather than representing sequential maturation stages. In addition, differential neuronal loss between PD and DLB may influence the cellular context in which pathology is sampled, such that the surviving neurons analysed here may represent distinct states of vulnerability rather than the full spectrum of disease-associated changes.

Furthermore, while our ultrastructural analysis identifies robust mitochondrial phenotypes, it does not resolve the underlying mechanisms driving these changes. The observed alterations may reflect a combination of impaired mitophagy, altered fusion–fission dynamics, and broader disruptions in cellular homeostasis. Establishing the biochemical and molecular drivers of these phenotypes will require complementary approaches, including spatial proteomics, lipidomics, and targeted analysis of mitochondrial quality control pathways.

Future studies incorporating larger cohorts, earlier disease stages, and volumetric or longitudinal approaches will be essential to define how mitochondrial phenotypes evolve over time and across regions. Integration of high-resolution structural methods (e.g. cryo-EM), and whole-cell or tissue-scale morphometrics will further enable comprehensive mapping of organelle-level pathology. Together, these approaches will help determine whether mitochondrial increased mitochondrial size and density represent a discriminating feature of PD or part of a broader organelle-wide dysfunction that emerges selectively in vulnerable neuronal populations.

## Conclusion

Together, these findings provide an ultrastructural framework for Lewy pathology across the human brain in PD and DLB. We show that α-Syn inclusions adopt region-specific architectures determined by neuronal identity, rather than disease type, and that mitochondrial homeostasis may represent a discriminator of synucleinopathy subtype. This work emphasizes that selective vulnerability arises from the intersection of α-Syn aggregation, intrinsic cellular environments, and regional context, highlighting critical features that must be captured in human-relevant disease models for both PD and DLB.

## Declarations

### Ethics approval and consent to participate

All donors provided written informed consent for a brain autopsy and the use of the material and clinical information for research purposes. Detailed neuropathological and clinical information was made available, in compliance with local ethical and legal guidelines, and all protocols were approved by the Amsterdam UMC institutional review board. Demographic features and clinical symptoms were abstracted from the clinical files, including sex, age at symptom onset, age at death, disease duration, presence of dementia, core and supportive clinical features for Parkinson’s disease and Dementia with Lewy bodies. All protocols were approved by VU University Medical Center institutional ethics review board (reference number 2019/148). This study was approved by the Cantonal Ethics Committee (Canton de Vaud, Project-ID CER-VD 2025-01061).

### Consent for publication

All donors provided written informed consent for the use of the material and clinical information for publication. All protocols were approved byVrije University Medical Center institutional review board. Fig. 7 was created with BioRender.com with agreement number ZY294VGV08.

### Data availability

All data will be made available on the BioImage Archive following acceptance of the manuscript for publication.

### Code availability

All codes are available as specified on GitHub: https://github.com/LBEM-CH/mitoanalyzer

## Author contributions

NS, AJL, and HS designed the study. WvdB collected, dissected and fixed the brain tissue samples for CLEM and performed the neuropathological diagnosis of the donors. NS, MS, AMRS, DS, LvdH, EA, and AJL performed the CLEM experiments. JR assisted with TEM imaging. MSK and MW wrote codes and trained datasets for cellpose. NS, SK and DP performed the mitochondria segmentation, assisted by MDF. NS performed all statistical analysis. NS, LvdH, MDF and AJL prepared the figures. NS and AJL wrote the manuscript. All authors contributed to the analysis and interpretation of the data and approved the final version of the manuscript.

## Acknowledgements

We are grateful to the individuals who participated in the brain donation program and their families, making this study possible. We would like to thank all members of the Netherlands Brain Bank autopsy team for facilitating the collection of high-quality postmortem brain tissue for electron microscopy. We thank the staff at the Electron Microscopy Facility at the University of Lausanne for sample preparation and TEM assistance and maintenance. We thank the EPFL Center for Imaging for useful discussions regarding image processing. We also thank Christel Genoud and Anne-Laure Mahul-Mellier for useful discussions.

