## Supplementary information for "Correlative Ultrastructural Mapping of Lewy Pathology Reveals Regional Diversity in Parkinson’s and Dementia with Lewy bodies"

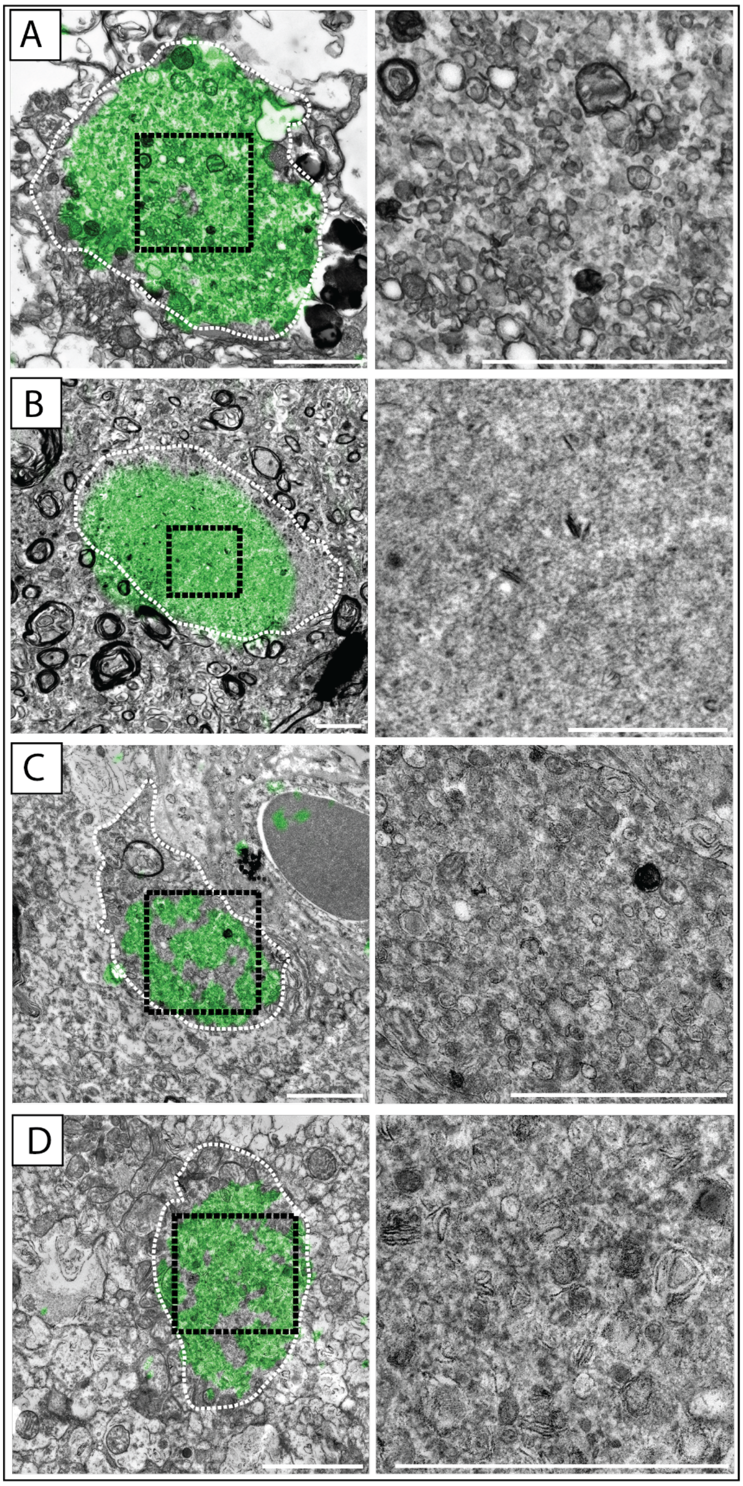


**Supplementary Figure 1. Neuritic pathology in ACC of DLB and PD cases.** In all panels, dashed lines outline neuritic boundaries; black boxed areas indicate higher magnification views. a) Membranous neuritic inclusion in the ACC of DLB case. b) Fully fibrillar neuritic inclusion in ACC of DLB case. c, d) Membranous neuritic inclusion in the ACC of PD cases. The IHC α-Syn immunostaining (green) is overlaid onto the EM micrograph. Scale bars: 2 µm.

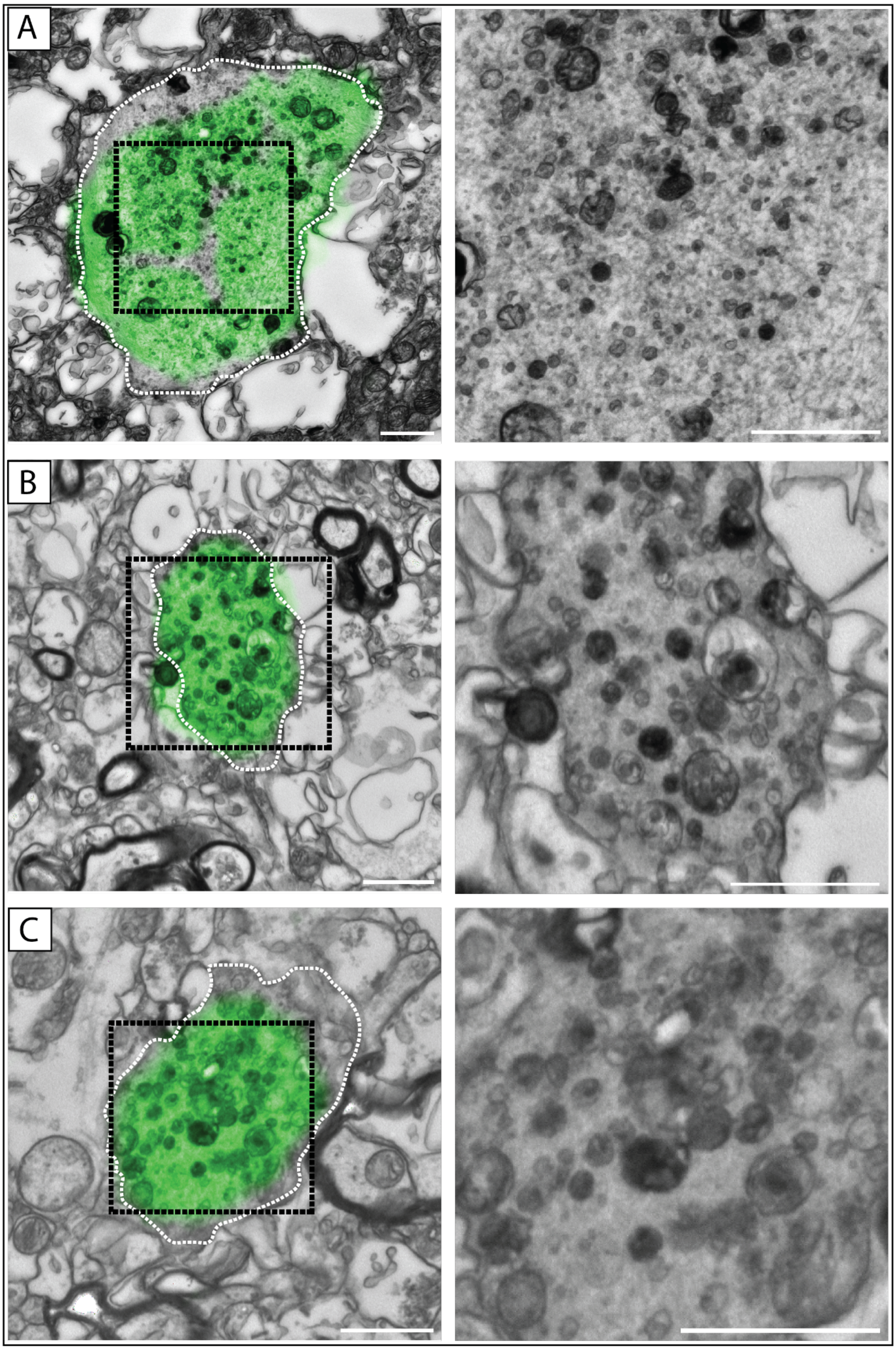


**Supplementary Figure 2. Neuritic pathology in ENT and CA2 of cases.** In all panels, dashed lines outline neuritic boundaries; black boxed areas indicate higher magnification views. a) Mixture of membranes and fibrils neuritic inclusion in the ENT of PD case. b) Mixture of membranes and fibrils neuritic inclusion in CA2 of PD case. c) Another example of mixture of membranes and fibrils neuritic inclusion in CA2 of PD case The IHC α-Syn immunostaining (green) is overlaid onto the EM micrograph. Scale bars: 1 μm.

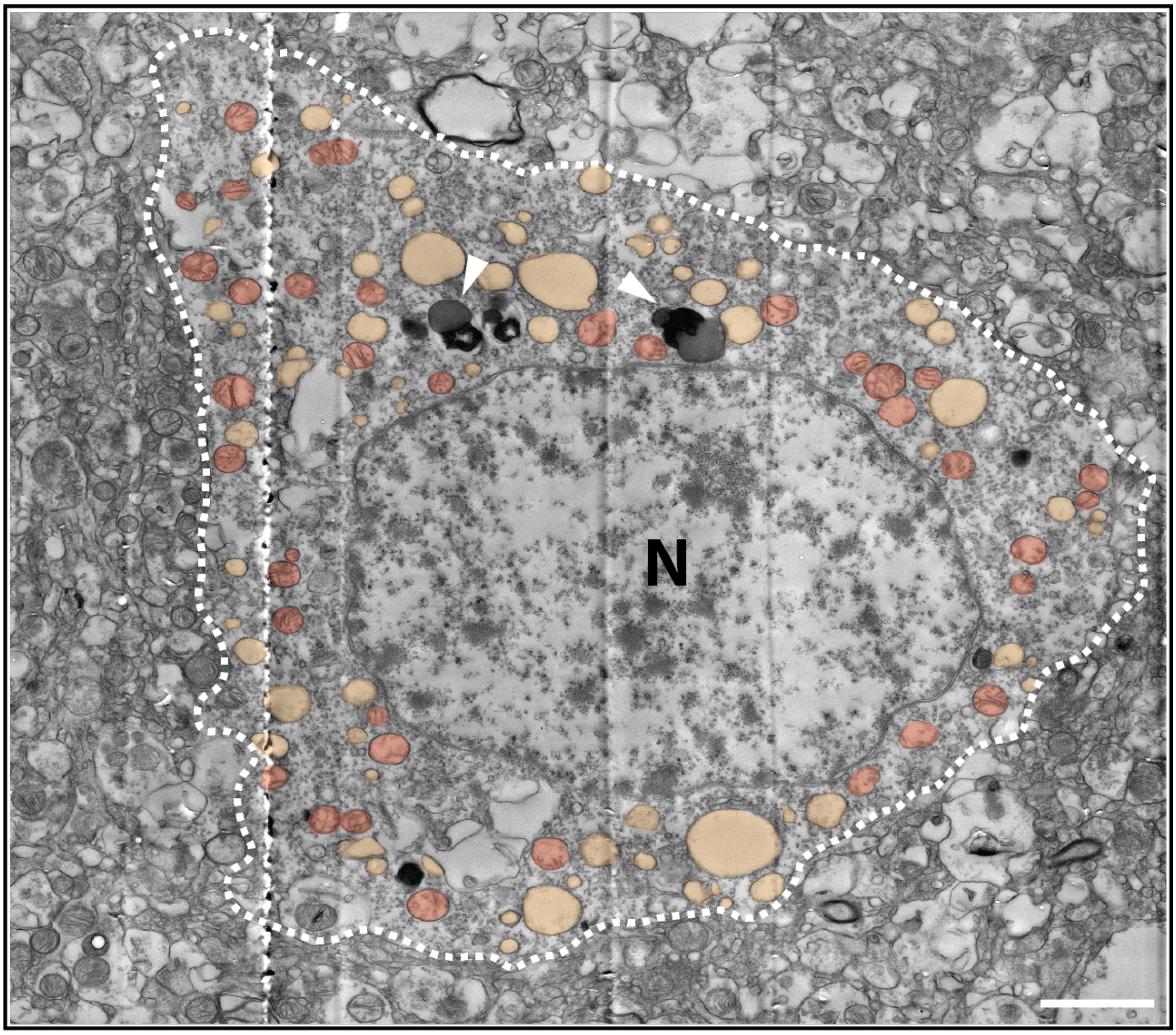


**Supplementary Figure 3. Neuron from the ENT of a non-neurological control.** N refers to the nucleus, and the membrane is indicated by the white dashed line. Mitochondria (red), other vesicles and organelles (yellow) are annotated. The nucleus is in the middle of the neuron, some lipofuscin droplets can be seen (white arrows), and there is no empty space between the neuron and the surrounding tissue. Scale bars: 2 µm

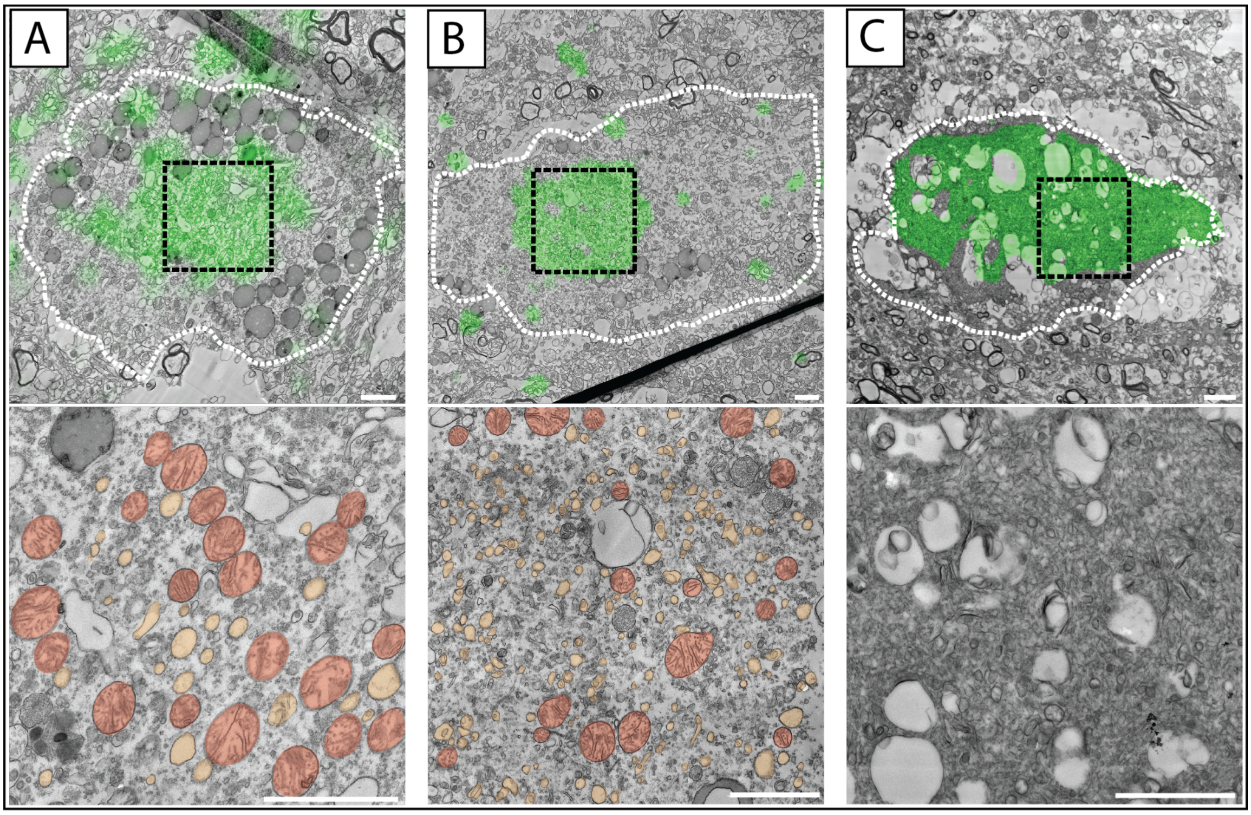


**Supplementary Figure 4. Irregular ultrastructures observed in the CA2 of DLB cases.** In all panels, dashed lines outline neuritic boundaries; black boxed areas indicate higher magnification views. Mitochondria (red), other vesicles and organelles (yellow) are annotated. a) The somatic ultrastructure in CA2, there are low density fibrils mixed with other organelles, b) Another example of somatic LB in CA2 with the ultrastructure as A. c) Irregular p-α-Syn positive region no fibrils have been observed. The IHC α-Syn immunostaining (green) is overlaid onto the EM micrograph. Scale bars: 2 µm.

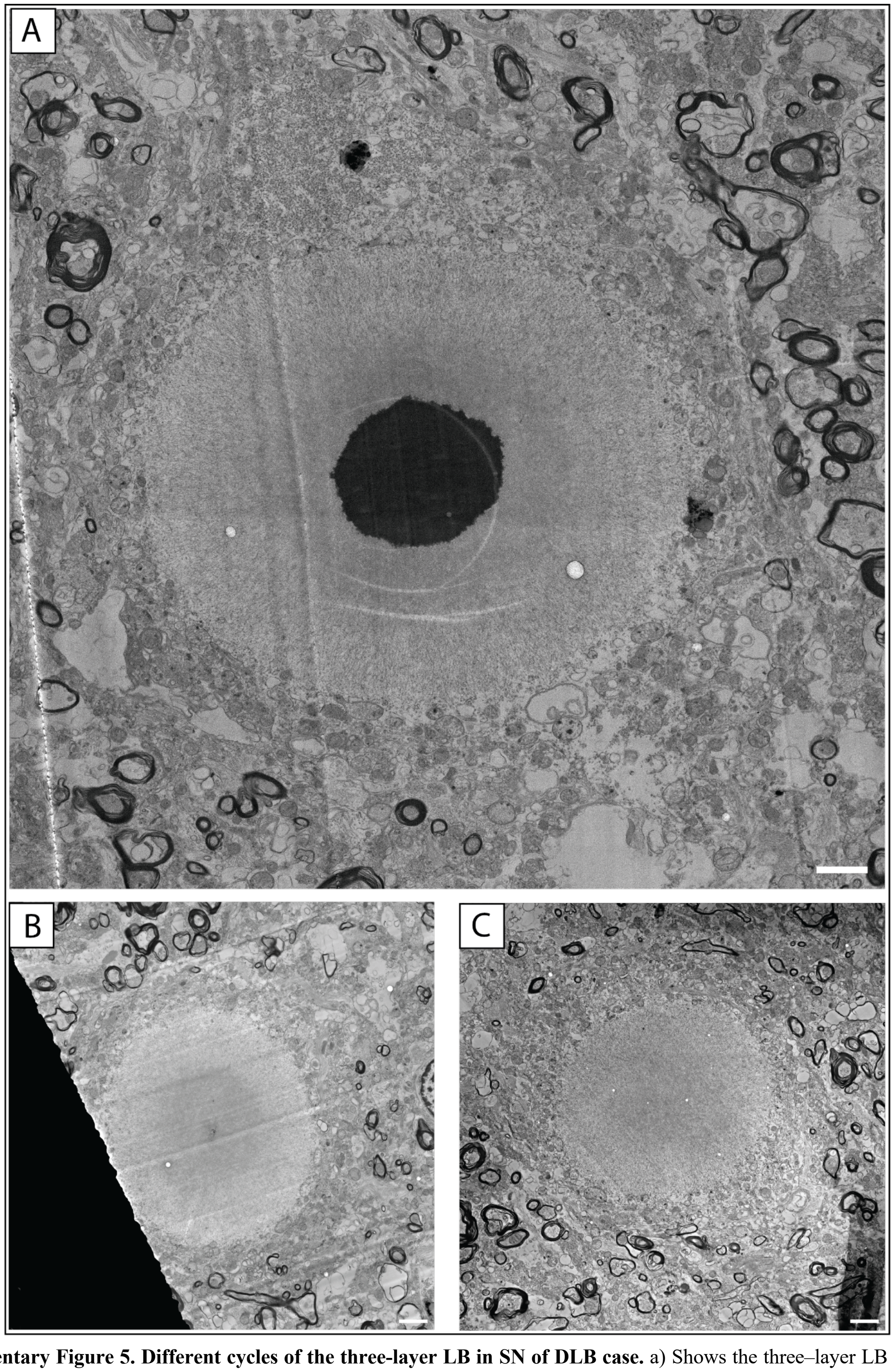


**Supplementary Figure 5. Different cycles of the three-layer LB in SN of DLB case.** a) Shows the three–layer LB in the first cycle. b) Shows the three–layer LB in third cycle. c) Shows the three–layer LB in the fourth cycle. The image shown in the main figure (Figure 3) obtained from cycle 2. Scale bars: 2 µm.

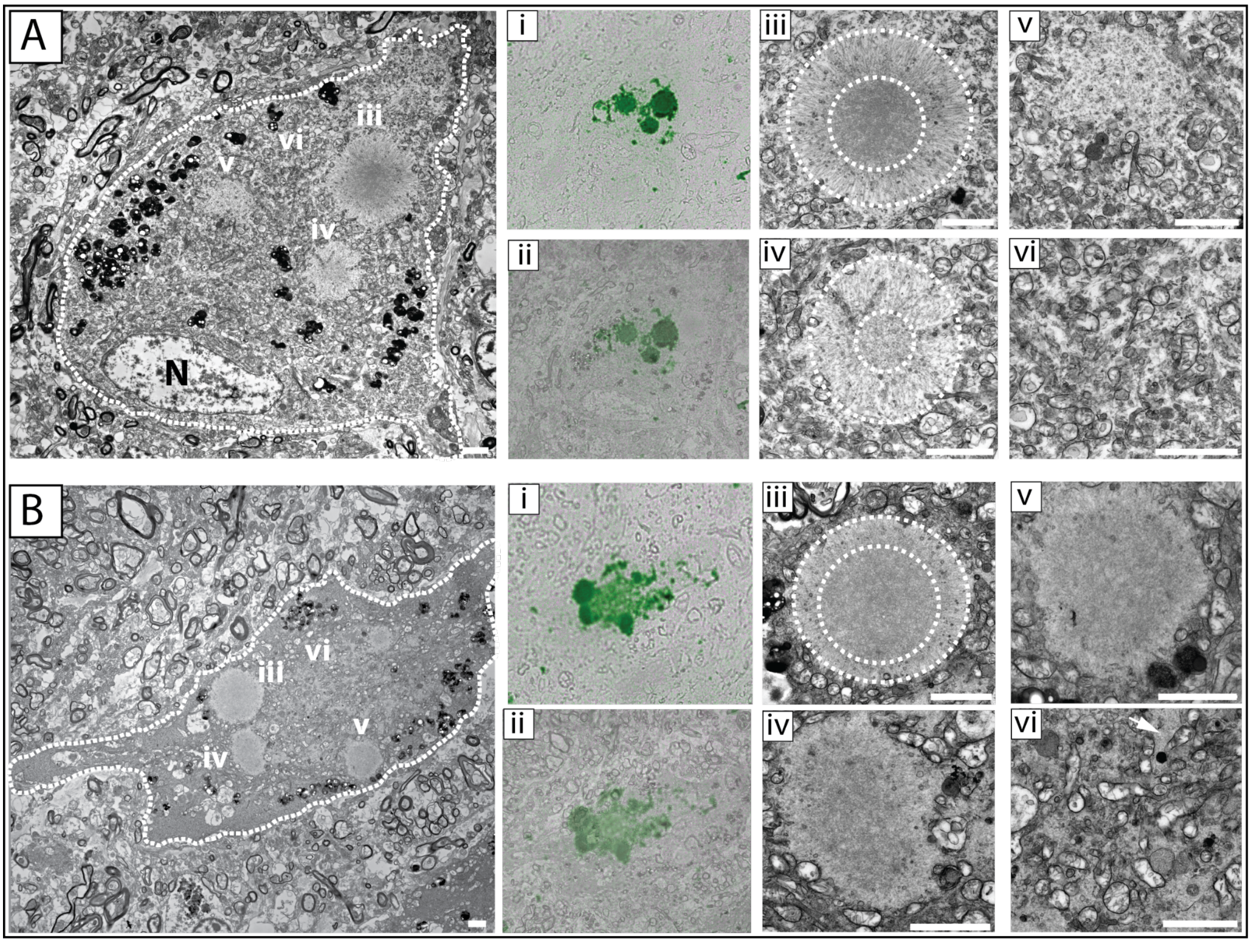


**Supplementary Figure 6. Examples of Multiple LBs in one neuron in SN of DLB cases.** a) Shows an example of multiple LB in one neuron which are connected by mitochondria. i) show α-Syn immunohistochemistry of the region, ii) shows the overlay of the EM and immunohistochemistry, iii) shows two-layer LB ultrastructure, iv) shows another two-layer LB as the second baby LB, v) shows one-layer LB which has no arranged fibrils, vi) show the area between these pathology which contains packed mitochondria with a few fibrils. b) Shows another example of multiple LBs in one neuron. i) show α-Syn immunohistochemistry of the region, ii) shows the overlay of the EM and immunohistochemistry, iii) shows two-layer LB ultrastructure, iv) shows one-layer LB, v) shows another one-layer LB, vi) show the pale body which connects LBs. Scale bars: 2 µm.

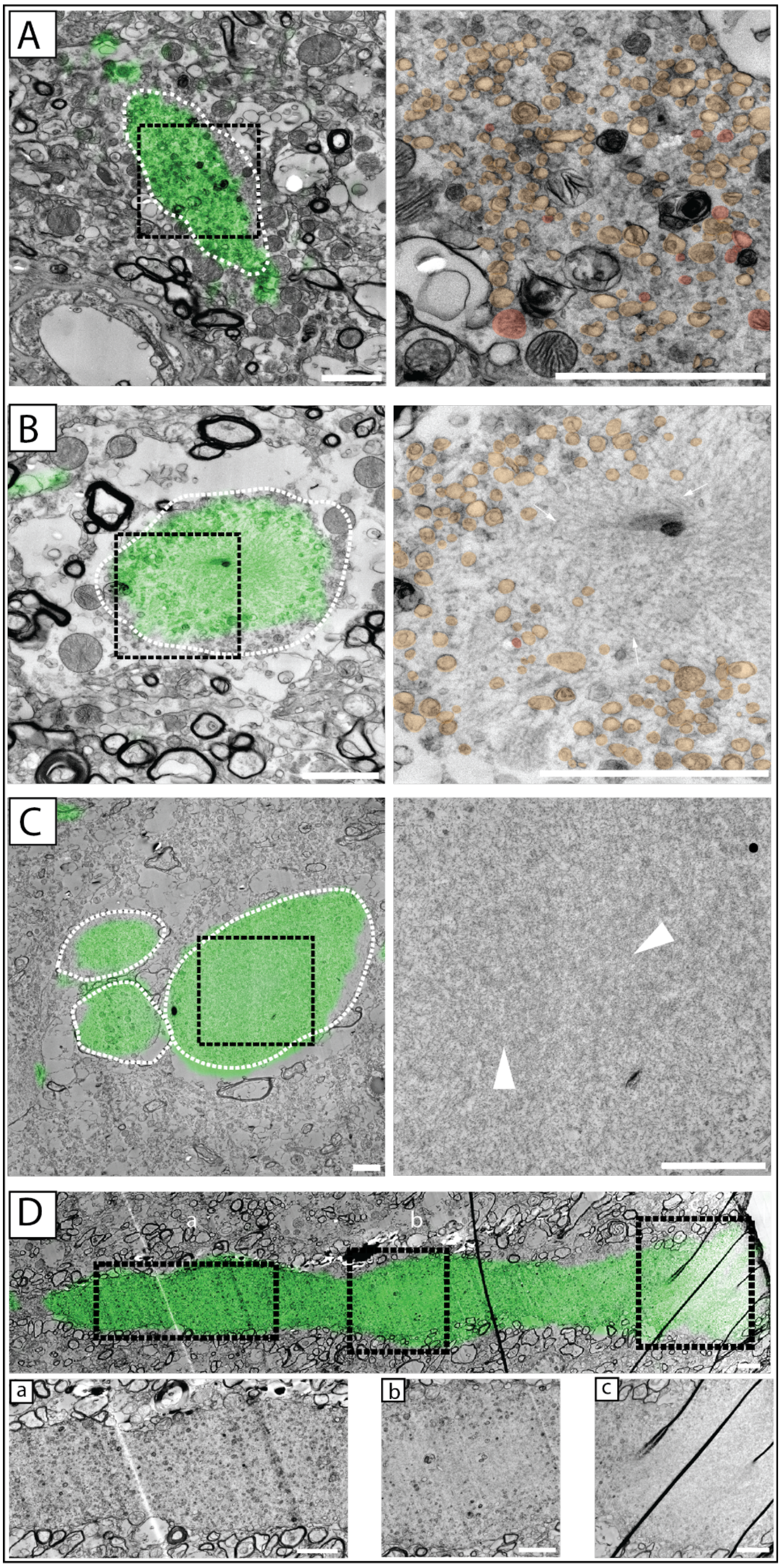


**Supplementary Figure 7. Neuritic pathology in CA2 in DLB cases.** In all panels, dashed lines outline neuritic boundaries; black boxed areas indicate higher magnification views. Mitochondria (red), other vesicles and organelles (yellow) and regions of fibrils (white arrowheads) are annotated. a) Membranous neuritic inclusion in the CA2. b) Central fibrillar and surrounded by other organelles. c) Fully fibrillar non somatic. d) Bulky neurite around 60 µm in length from the CA2 (top original image indicating cut-outs, bottom with IHC overlaid). Cutouts in i) show mixed fibrillar and other organelles, ii) more fibrillar, and iii) fully fibrillar but not positive for p-α-Syn. The IHC α-Syn immunostaining (green) is overlaid onto the EM micrograph. Scale bars: 2 µm.

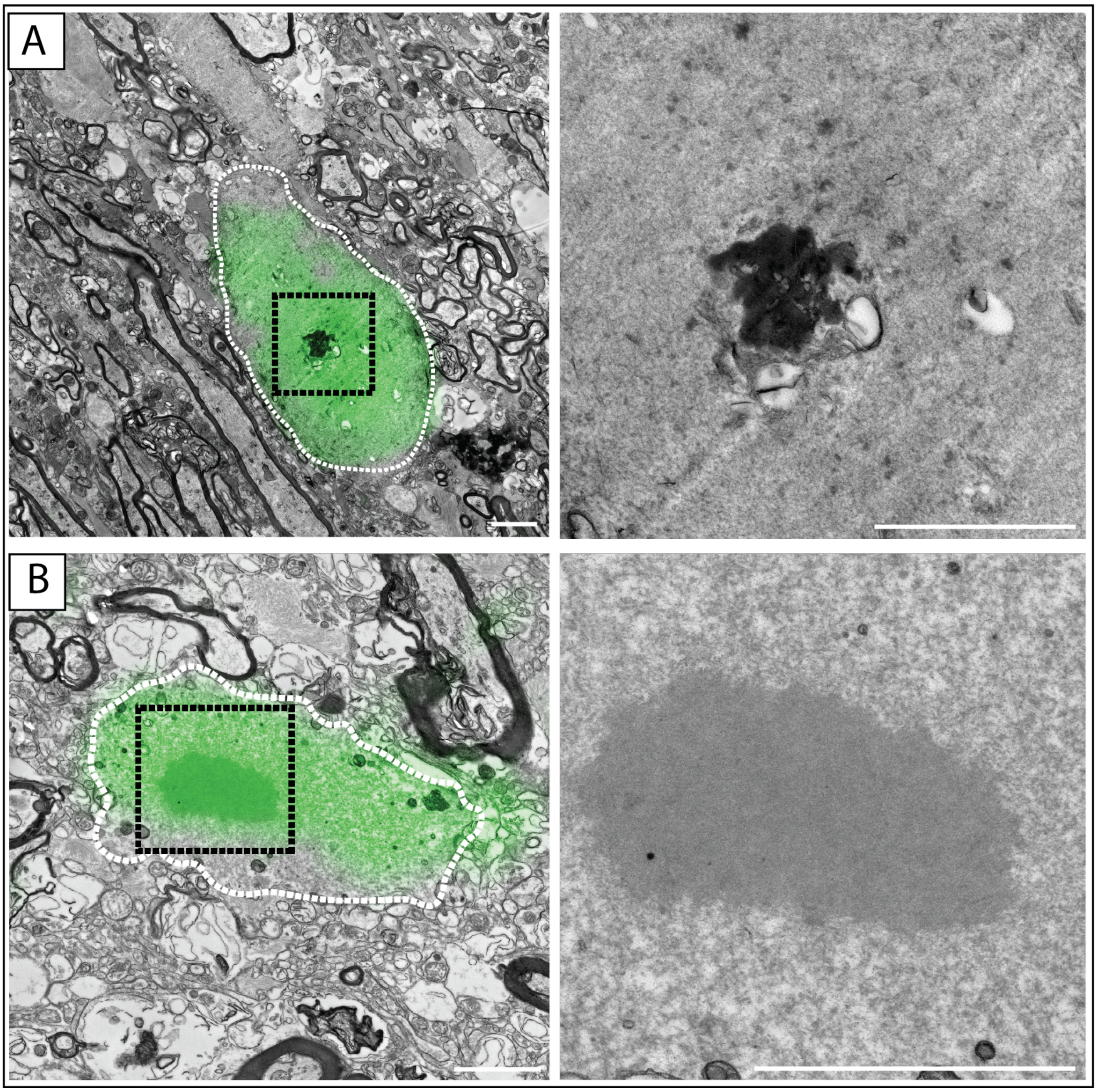


**Supplementary Figure 8. Other examples of neuritic pathology with electron dense core in SN of DLB cases.** In all panels, dashed lines outline neuritic boundaries; black boxed areas indicate higher magnification views. a) shows the example of fibrillar neuritic pathology with a dark core. b) Shows another example with lower electron core density**.** The IHC α-Syn immunostaining (green) is overlaid onto the EM micrograph. Scale bars: 2 µm.

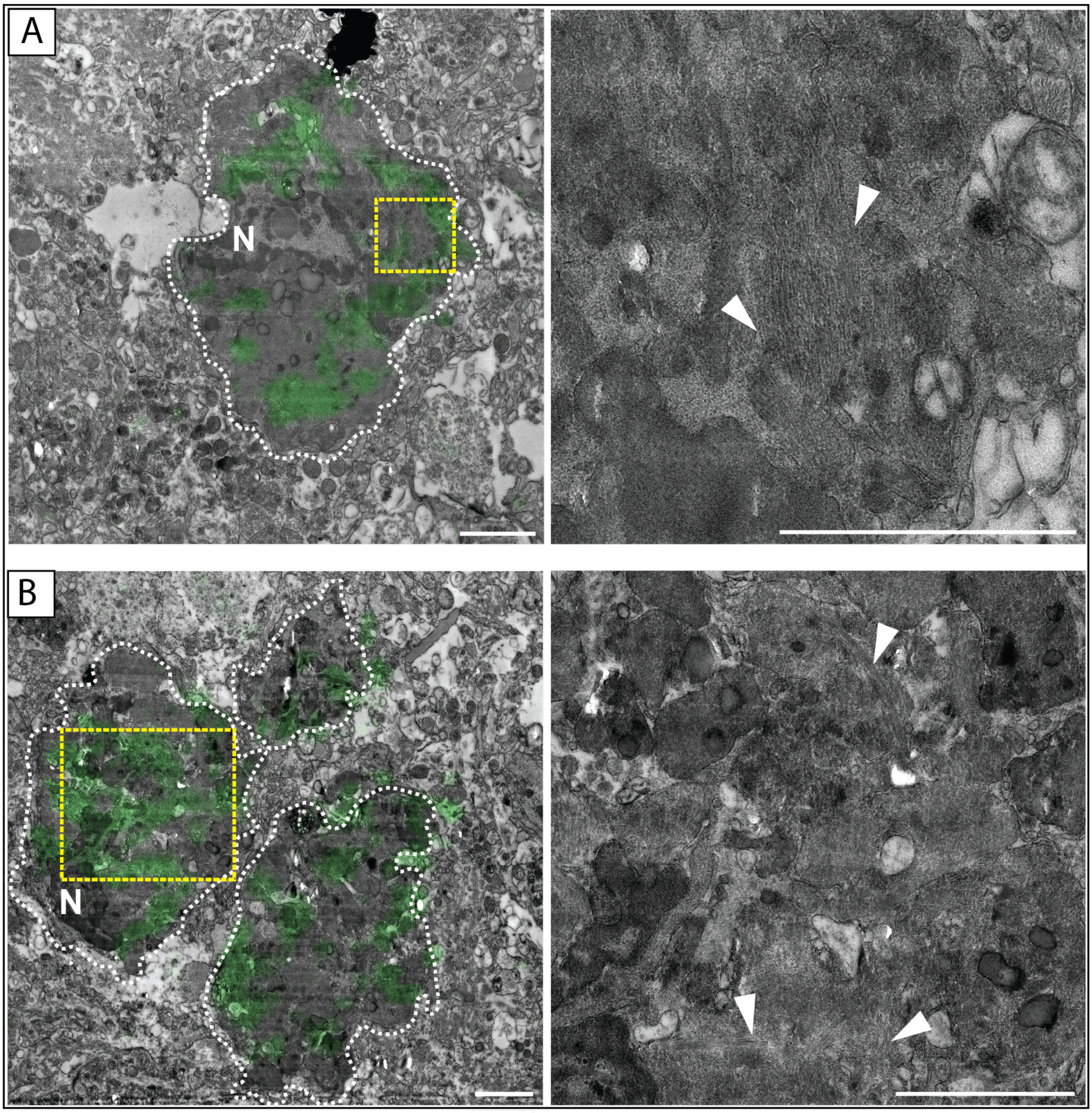


**Supplementary Figure 9. Irregular pathology in ENT from DLB case.** In all panels, dashed lines outline neuritic boundaries; yellow boxed areas indicate higher magnification views. Regions of fibrils (white arrowheads) are annotated. a) Degenerating neuron positive for p-α-Syn in the ENT; the nucleus appears dark. b) Somatic and non-somatic α-Syn-positive region associated with the degenerating neuron. Dense packed oriented fibrils are indicated by white arrow. The IHC α-Syn immunostaining (green) is overlaid onto the EM micrograph. Scale bars: 2 μm.

| Case | F/M | Age at death (y) | Diagnosis | | PMD (h:m) | Brain weight (g) | Braak α-Syn stage | Braak NFT | Thal phase |
|---|---|---|---|---|---|---|---|---|---|
| | | | Clin. | Path. | | | | | |
| DLB 1 | M | 72 | DLB | DLB+AD | 04:15 | 1397 | 6 | 3 | 4 |
| DLB 2 | M | 71 | DLB | DLB | 04:15 | 1400 | 6 | 2 | 2 |
| DLB 3 | M | 68 | DLB | DLB | 04:55 | 1605 | 6 | 2 | 3 |
| DLB 4 | F | 77 | DLB | DLB | 06:45 | 1185 | 6 | 2 | 3 |
| PD 1 | M | 88 | PDD | PDD | 4:05 | 1405 | 6 | 3 | 3 |
| PD 2 | M | 82 | PDD | PDD | 6:50 | 1200 | 6 | 2 | 0 |
| PD 3 | M | 83 | PD | PD | 6:25 | 1387 | 6 | 3 | 1 |
| PD 4 | M | 74 | PD | PDD | 5:15 | 1350 | 6 | 2 | 3 |
| PD 5 | M | 84 | PD | PD | 4:50 | 1430 | 5 | 3 | 0 |
| Cont 1 | F | 74 | Cont | Cont | 6:10 | n.d | 1 | 2 | 3 |
| Cont 2 | M | 63 | Cont | Cont | 3:00 | 1350 | 0 | 1 | 1 |

**Supplementary Table 1**. Clinicopathological features of donors included in this study. DLB=Dementia with Lewy bodies; PD=Parkinson's disease; AD=Alzheimer disease; Cont=control; F/M=female/male; y=year; Clin.=clinical diagnosis; Path.=pathological diagnosis; PMD= postmortem delay; h=hours; m=minutes; g=grams;NFT=neurofibrillary tangle. n.d= no data available.

| Disease | Brain region | Fibrillar density | | | Total # inclusions | |
|---|---|---|---|---|---|---|
| | | Low density | Mid density | High density | | |
| Cortical LBs | | | | | Per brain region | Per disease |
| DLB | ENT | 19 | 20 | 2 | 41 | 67 |
| | ACC | n.o. | n.o. | 22 | 22 | |
| | CA2 | 2 | n.o. | n.o. | 2 | |
| PD | ENT | 9 | 4 | 3 | 16 | 26 |
| | ACC | n.o. | n.o. | 10 | 10 | |
| Degenerating neurons | | | | | | |
| DLB | ENT | 2 | 3 | n.o. | 5 | 7 |
| | ACC | n.o. | 0 | 2 | 2 | |
| PD | ACC | n.o. | n.o. | 1 | 1 | 1 |

**Supplementary Table 2. Classification and count of cortical α-Syn pathology localized by CLEM.** DLB=Dementia with Lewy bodies; PD=Parkinson's disease; ENT=entorhinal cortex; ACC= anterior cingulate cortex; CA2=hippocampus; n.o. = not observed.
